# EPSD 2.0: An Updated Database of Protein Phosphorylation Sites across Eukaryotic Species

**DOI:** 10.1101/2025.01.13.632887

**Authors:** Miaomiao Chen, Yujie Gou, Ming Lei, Leming Xiao, Miaoying Zhao, Xinhe Huang, Dan Liu, Zihao Feng, Di Peng, Yu Xue

## Abstract

As one of the most crucial post-translational modifications (PTMs), protein phosphorylation regulates a broad range of biological processes in eukaryotes. Biocuration, integration and annotation of reported phosphorylation events will deliver a valuable resource for the community. Here, we present an updated database, the eukaryotic phosphorylation site database 2.0 (EPSD 2.0), which includes 2,769,163 experimentally identified phosphorylation sites (p-sites) in 362,707 phosphoproteins from 223 eukaryotes. From the literature, 873,718 new p-sites identified through high-throughput phosphoproteomic research were first collected, and 1,078,888 original phosphopeptides together with primary references were reserved. Then, this dataset was merged into EPSD 1.0, comprising 1,616,804 p-sites within 209,326 proteins across 68 eukaryotic organisms [1]. We also integrated 362,190 additional known p-sites from 10 public databases. After redundancy clearance, we manually re-checked each p-site and annotated 88,074 functional events for 32,762 p-sites, covering 58 types of downstream effects on phosphoproteins, and regulatory impacts on 107 biological processes. In addition, phosphoproteins and p-sites in 8 model organisms were meticulously annotated utilizing information supplied by 100 external platforms encompassing 15 areas. These areas included kinase/phosphatase, transcription regulators, three-dimensional structures, physicochemical characteristics, genomic variations, functional descriptions, protein domains, molecular interactions, drug-target associations, disease-related data, orthologs, transcript expression levels, proteomics, subcellular localization, and regulatory pathways. We expect that EPSD 2.0 will become a useful database supporting comprehensive studies on phosphorylation in eukaryotes. The EPSD 2.0 database is freely accessible online at https://epsd.biocuckoo.cn/.

## Introduction

Protein phosphorylation serves as a crucial post-translational modification (PTM), orchestrating a wide range of biological functions [2–5]. In eukaryotes, phosphorylation primarily modifies specific amino acids, especially serine (S), threonine (T), or tyrosine (Y) residues on targeted substrates catalyzed by protein kinases [6]. Phosphorylation at different sites on a protein may exert distinct functions under various biological contexts [2]. For example, phosphorylation of the human PLK1 protein at S137 promotes cancer cell survival in response to mitochondrial dysfunction [7], while the phosphorylation at T210 is required for enhancing neuroprotective autophagy [8]. In particular, aberrant or inadequate phosphorylation has been implicated in multiple diseases, including neurodegenerative diseases [9], cancer [10], and cardiovascular disorders [11]. Consequently, pinpointing phosphorylation sites (p-sites) in eukaryotes is essential for understanding the regulatory patterns and underlying mechanisms of phosphorylation in diverse biological processes.

Tandem mass spectrometry (MS/MS) serve as a leading technique for identifying p-sites in proteins [12,13]. Besides Orbitrap mass spectrometers [14], recent progress has led to the development of MS/MS instruments with much higher speed, throughput, and resolution. For example, by incorporating ion mobility spectrometry as a new dimension, MS/MS technology has advanced to a four-dimensional (4D) stage [15,16]. Utilizing trapped ion mobility spectrometry coupled with a time-of-flight (timsTOF) mass spectrometer, Kramer et al. identified nearly 30,000 p-sites from bone marrow samples of 44 acute myeloid leukemia patients and 6 healthy controls [17]. Recently, the Asymmetric Track Lossless (Astral), a novel type of mass analyzer, achieved higher scanning speed, resolving power, sensitivity, and low-ppm mass accuracy [18,19]. Coupled with Orbitrap, the leading Orbitrap Astral mass spectrometer helped in identifying 81,120 unique p-sites within 12 hours of measurement in a data-independent acquisition (DIA) manner [20]. These advancements in MS/MS have facilitated the continuous identification of numerous p-sites.

To manage and utilize these data, many public resources have been developed to curate p-sites, including Phospho.ELM [21,22], PhosphoSitePlus [23,24], dbPTM [25,26], SysPTM [27,28], iPTMnet [29,30], PHOSIDA [31], PhosphoPep [32], LymPHOS [33,34], PlantsP [35], PhosPhAt 4.0 [36,37], P3DB [38], MPPD [39], FPD [40], UniProt [41,42], Plant PTM Viewer [43], HPRD [44], BioGRID [45], RegPhos [46,47], Pf-phospho [48], Scop3P [49], HisPhosSite [50], and Nphos [51] (Supplementary Table S1). Among these databases, PhosphoSitePlus, dbPTM, SysPTM, iPTMnet, Plant PTM Viewer, and UniProt provide data for multiple PTM types in addition to phosphorylation. PhosphoSitePlus and dbPTM also offer functional annotations on the downstream effects of p-sites. Additionally, most of the existing resources mainly curate *O*-phosphosites, whereas HisPhosSite and Nphos focus on the collection of *N*-phosphosites. Of note, Nphos contains 11,710 experimentally verified *N*-phosphosites covering pHis, pLys, and pArg residues, and provides an online service for predicting *N*-phosphosites, serving as a highly useful resource for *N*-phosphorylation [51]. With the development of artificial intelligence (AI) technologies, more structured and comprehensive datasets are needed to support the training of large models. Therefore, there remains a need for the collection, curation and combination for extensive constantly experimentally verified p-sites. Also, it is very desirable to obtain the specific functional mechanisms of phosphorylation events, so it is important and useful to annotate the downstream effects of p-sites.

In 2019, we released EPSD 1.0 that comprised 1,616,804 p-sites by re-curating data from the dbPPT [52] and dbPAF [53] databases, along with p-sites identified in high-throughput (HTP) studies and 13 publicly accessible phosphorylation databases. In this release of EPSD 2.0, we further re-curated the p-sites in EPSD 1.0, collected p-sites from HTP experiments and 10 additional public databases (Figure 1A). This resulted in a dataset comprising 2,769,163 p-sites on 362,707 phosphoproteins across 223 species, encompassing 3,364,760 phosphopeptides. Furthermore, we annotated 88,074 functional events for 32,762 p-sites through extensive literature review. These annotations of downstream effects covered 58 types of functions on phosphoproteins, and 107 regulatory impacts on biological processes. Additionally, we meticulously annotated the dataset by incorporating the information from 100 extra resources. Ultimately, EPSD 2.0 (∼36.2 GB) exhibits roughly a 2.5-fold increase in the data volume compared to EPSD 1.0 (∼14.1 GB). Compared to other existing databases, EPSD 2.0 offers more experimentally validated p-sites across more eukaryotic species, alongside extensive functional annotations on downstream effects of p-sites. We believe that EPSD 2.0 will serve as a more comprehensive and valuable database for the scientific community.

**Figure 1.**
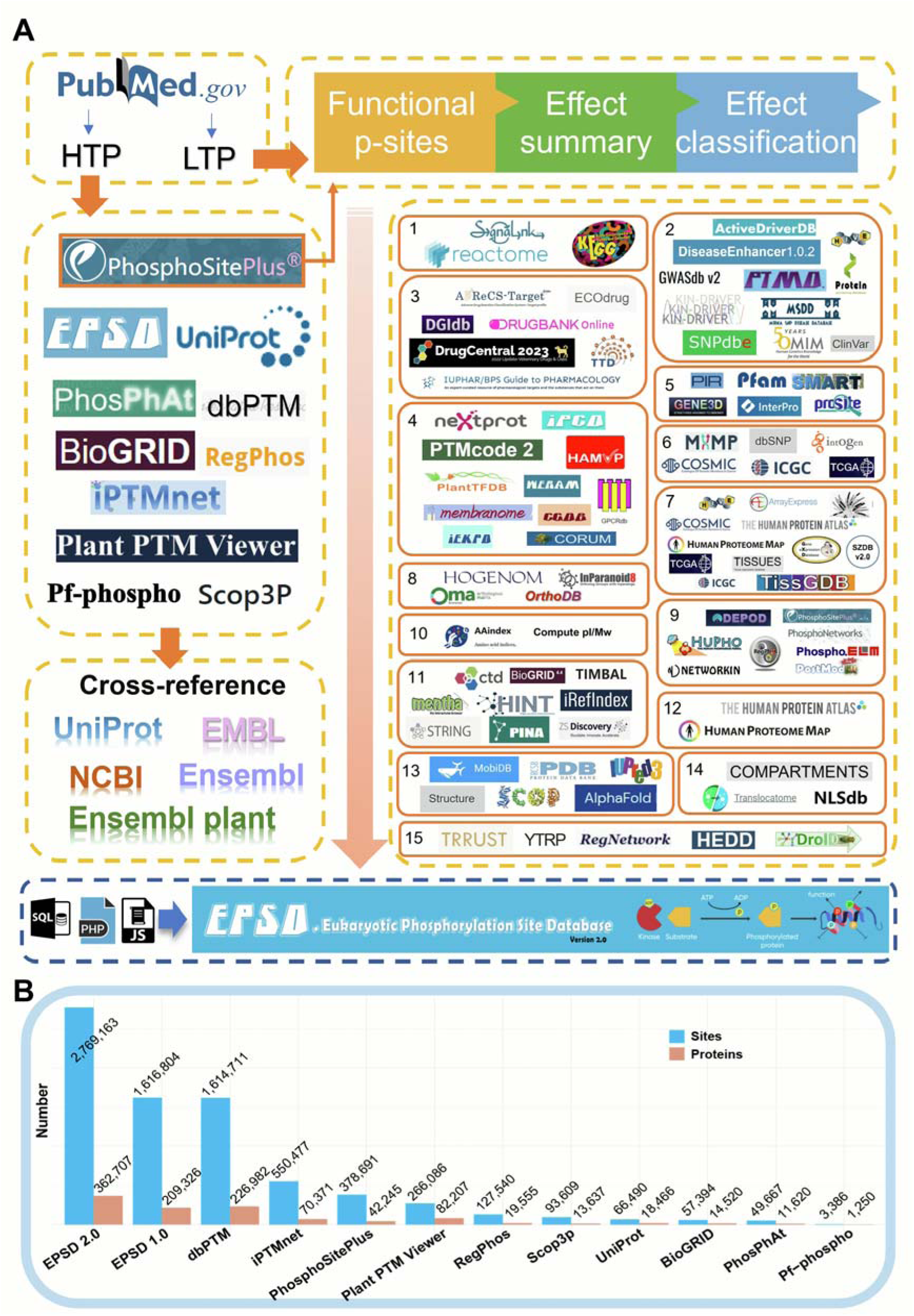
The procedure for development of EPSD 2.0. **A**. First, we manually collected experimentally identified p-sites from PubMed. Then, known p-sites in 10 additional phosphorylation databases were integrated. In addition to basic annotations, we further annotated eight model organisms, including *Homo sapiens*, *Mus musculus*, *Rattus norvegicus*, *Drosophila melanogaster*, *Caenorhabditis elegans*, *Arabidopsis thaliana*, *Schizosaccharomyces pombe* and *Saccharomyces cerevisiae*, by integrating the knowledge from 100 additional resources that covered 15 aspects, including kinase/phosphatase, transcription regulators, three-dimensional structures, physicochemical characteristics, genomic variations, functional descriptions, protein domains, molecular interactions, drug-target associations, disease-related data, orthologs, transcript expression levels, proteomics, subcellular localization, and regulatory pathways. **B**. The numbers of p-sites and phosphoproteins curated and integrated from EPSD and other existing resources.

### Construction and content

#### Data collection, curation and integration

This update not only expanded the collection of new p-sites but also curated the downstream effects of these p-sites (Figure 1). First, we performed a PubMed search employing various keywords, such as “large-scale phosphorylation”, “phosphoproteome”, “phosphoproteomics”, “phosphoproteomic”, “mass spectrometry phosphorylation”, “MS phosphorylation”, “phospho-residue”, “phospho-site”, and “phosphosite”. This search aimed to get published studies containing HTP phosphoproteomic data. In these HTP studies, different MS/MS instruments were used for phosphoproteomic profiling, and different software packages were used for data processing. However, all these studies followed a standard procedure for data processing, in which the false discovery rates for the peptide-spectrum match, p-site and protein decoy fraction were all set to < 1%, supporting the high quality of all studies. To ensure the consistency with the original studies, we directly obtained HTP p-sites from the supplementary data of each paper if available. Additionally, to capture articles containing p-sites with descriptions of their downstream effects, we conducted targeted searches on PubMed using combinations of keywords such as “phosphorylation”, “phosphorylated”, “phospho-” along with “site”, “residue”, “Serine”, “Threonine”, “Tyrosine”, “Ser”, “Thr”, and “Tyr”. Moreover, we reviewed the literature that served as the source of existing low-throughput (LTP) p-sites in the EPSD 2.0 database, aiming to extract descriptions of functional information related to p-sites. Here, we defined a functional event as a specific p-site coupled with one of its reported downstream effects. To ensure the data quality, the primary references were presented for both HTP and LTP p-sites.

To obtain the position information, all the modified residues with collected peptides were aligned with the reference sequences obtained from the UniProt database [42]. For the annotation of fundamental information of modified proteins, we also retained relative data from UniProt [42], including protein names, gene names, gene synonyms, Ensembl IDs, functional information, keywords, and sequences. In total, we obtained 873,718 nonredundant new p-sites of 125,180 proteins from 575 HTP phosphoproteomic studies (Figure 1B). By collecting functional p-sites with their summarized descriptions from PubMed and PhosphositePlus [24], we classified the functional descriptions into short categories, and finally retained 88,074 functional downstream events for 32,762 p-sites.

Next, these p-sites were merged with the existing dataset in EPSD 1.0 [1] and ten other public databases, including PhosphoSitePlus [24], dbPTM [26], UniProt [42], PhosPhAt [37], BioGRID [45], RegPhos [47], iPTMnet [30], Plant PTM Viewer [43], Pf-phospho [48], and Scop3P [49] (Figure 1A). For each database, the obtained phosphopeptides were remapped to the benchmark sequences of UniProt, and exact positions of p-sites were pinpointed. To ensure the consistency and reliability of data, the origins of p-sites obtained from other databases were documented. Ultimately, we obtained 2,769,163 nonredundant p-sites on 362,707 phosphoproteins across 223 species with annotations of 88,074 functional events (Figure 1B, Supplementary Table S3, S4). In comparison to EPSD 1.0 and other phosphorylation databases, EPSD 2.0 exhibited an approximate 71% enhancement in collection of p-sites and a nearly 2-fold increase in functional events.

#### Multilayer annotation for phosphoproteins and phosphorylation sites

Designed as a protein-centered database, EPSD 2.0 assigned each phosphoprotein with an automatically generated EPSD ID (EP-) as the main identifier, and utilized a UniProt accession as the alternative identifier, e.g., EP0017523 corresponds to the PLK1 protein in *Homo sapiens* with UniProt ID P53350. To provide detailed information for each phosphoprotein entry, some basic annotations were integrated from UniProt [42], for example, accession numbers of UniProt/RefSeq/Ensembl, NCBI Taxa ID, gene name and synonyms, NCBI gene ID, protein name and synonyms, protein and nucleotide sequences, Gene Ontology (GO) terms, and keywords. Additionally, the representative three-dimensional (3D) structure of the protein is provided, if available, from the PDB [38] or AlphaFold [54] database, with p-sites visualized on the structure. For each p-site, a flanking peptide of 15 amino acids (aa) centered around the middle-phosphorylated residue was provided with its reference information and PubMed IDs. For p-sites from HTP experiments, the initial peptides detected via MS and their cell or tissue origins were preserved when available. MS data analyzing software like MaxQuant can compute and assign the localization probability (LP) score for every candidate p-site [55]. The LP score quantifies the likelihood of correctly identifying phosphorylation sites within protein sequences based on mass spectrometry data. The LP score varies between 0 and 1, with higher values indicating a greater probability that a site is an authentic p-site. Accordingly, we classified the p-sites with pre-calculated LP scores directly obtained from these studies into four classes: I (> 0.75), II (0.5–0.75), III (0.25–0.5) and IV (< 0.25) [55]. To ensure the consistency with the original papers, we retained all p-sites from HTP experiments without applying any additional filters, following the settings of EPSD 1.0.

In addition to the basic information, we reviewed each functional description from literatures, summarized them into short phrases, and classified them into two types: “Effect on Protein” and “Effect on Biological Process” (Figure 1A). For example, “enzymatic activity, induce” falls under “Effect on Protein”, while “cell growth, inhibit” is one category of “Effect on Biological Process”. Moreover, we meticulously annotated phosphoproteins in eight model organisms, leveraging information from 100 external resources (Figure 1A). The annotation covered 15 different aspects, including kinase/phosphatase, transcription regulators, 3D structures, physicochemical characteristics, genomic variations, functional descriptions, protein domains, molecular interactions, drug-target associations, disease-related data, orthologs, transcript expression levels, proteomics, subcellular localization, and regulatory pathways. (Figure 1A, Supplementary Table S2). These annotations were directly used, and no inconsistencies were found. The Supplementary Methods (Supplementary Text) provided in-depth details on how annotations from each resource were processed. The p-sites and annotation datasets in EPSD 2.0 are available at https://epsd.biocuckoo.cn/Download.php.

#### The data statistics in the EPSD 2.0

Compared to EPSD 1.0, the updated EPSD 2.0 has added 1,152,359 p-sites and 153,381 new phosphoproteins. Additionally, the number of eukaryotes has increased by over 3-fold to 223 species, including 95 animals, 20 protists, 61 plants, 48 fungi. The heatmap in Figure 2A illustrates the number of phosphoproteins, p-sites, phospho-serine (pS), phospho-threonine (pT), and phospho-tyrosine (pY) residues across the top 80 organisms. The specific numbers for all organisms were presented in Supplementary Table S3. The results revealed that *H. sapiens* and *Mus musculus* occupy approximately 57.99% of all the p-sites.

**Figure 2.**
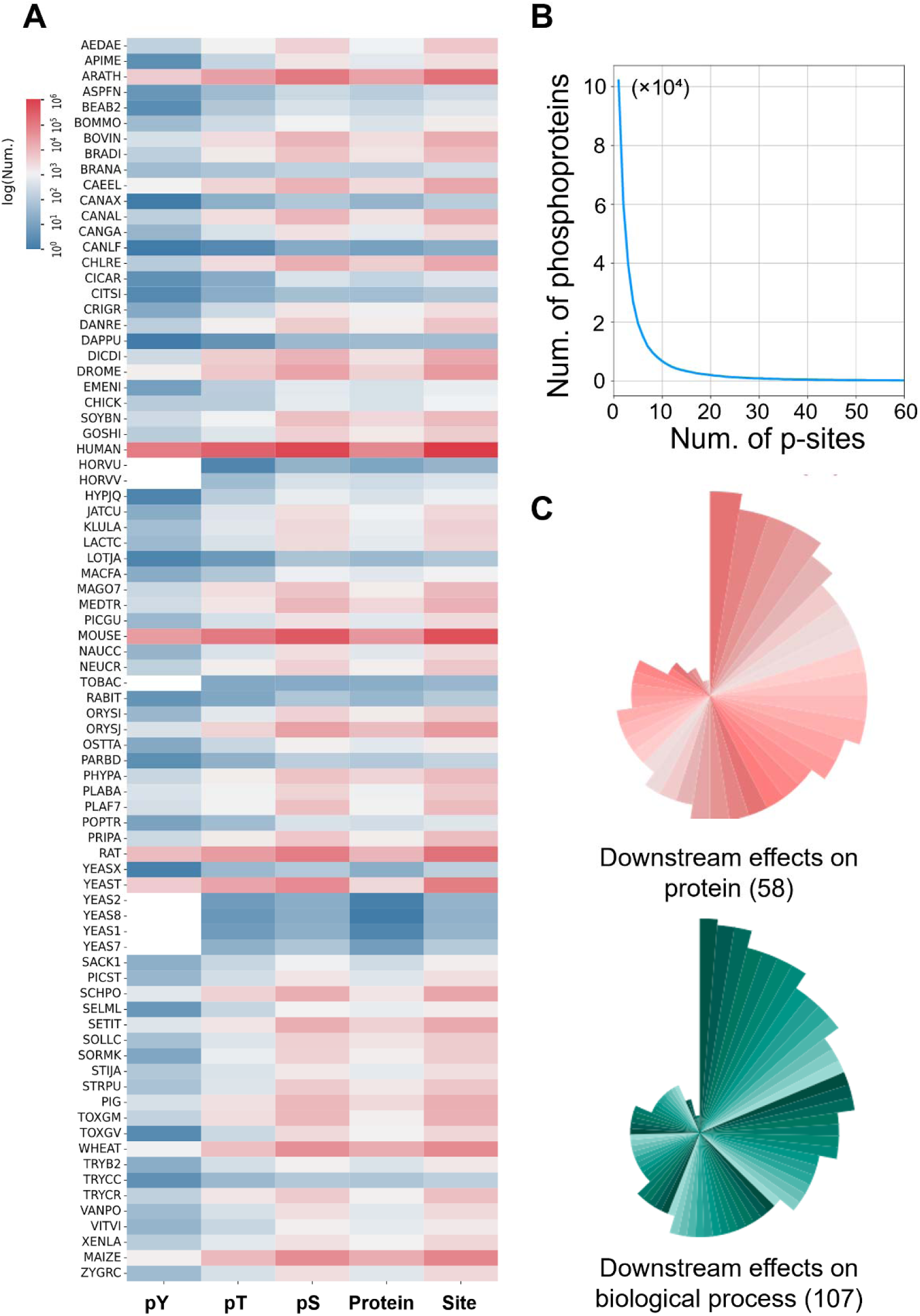
The data statistics in EPSD 2.0. **A**. A heatmap of the distribution of p-sites, phosphoproteins, and residues for each species in EPSD 2.0. **B**. The distribution of numbers of p-sites in protein substrates. **C**. The downstream effects on protein function and biological process.

EPSD 1.0 did not provide descriptions of potential functional effects related to phosphorylated sites. In EPSD 2.0, we annotated 88,074 functional downstream events on 32,762 p-sites, including 58 categories related to protein function and 107 categories about biological processes (Figure 2C, 2D). Notably, the most common effect on proteins is “molecular association, regulation”, indicating that phosphorylation on proteins usually alters the interactions with other molecules. Additionally, when a site was phosphorylated, its downstream effects on biological processes most frequently involved in the disease progression.

Among all the p-sites, there were 1,930,151 (69.70%) pS, 638,944 (23.07%) pT and 200,020 (7.22%) pY residues, with the remaining p-sites involving other residues like histidine. The distribution of p-sites per phosphoprotein revealed that 260,683 phosphoproteins (71.87%) were phosphorylated with at least two p-sites, highlighting multisite phosphorylation as a dominant regulatory mechanism for substrate phosphoproteins (Figure 2B). Specifically, 7840 (2.16%) phosphoproteins exhibited 50 or more p-sites, demonstrating the highly complex phosphorylation regulatory patterns in these proteins (Figure 2B).

#### Usage

EPSD 2.0 features an intuitive and easy-to-navigate interface. Using the human PLK1 protein, we demonstrate how to utilize the EPSD 2.0 website. Users can perform a one-click search on the home or search page by selecting “Protein Name” as the search option and entering the corresponding name “Serine/threonine-protein kinase PLK1” in the input box of “Substrate Search” field (Figure 3A). After clicking the “Submit” button or pressing “Enter”, a table will be generated to display the search results, which includes the columns of EPSD ID, UniProt Accession, Gene Name, Protein Name, and Species (Figure 3A). The same result can also be obtained from the browse page by clicking the “*Homo sapiens*” in the “Browse by Species” section (Figure 3A). Then, a “View” page will be displayed to provide more detailed information by clicking on “EP0017523”, the EPSD ID corresponding to the human PLK1 protein.

**Figure 3.**
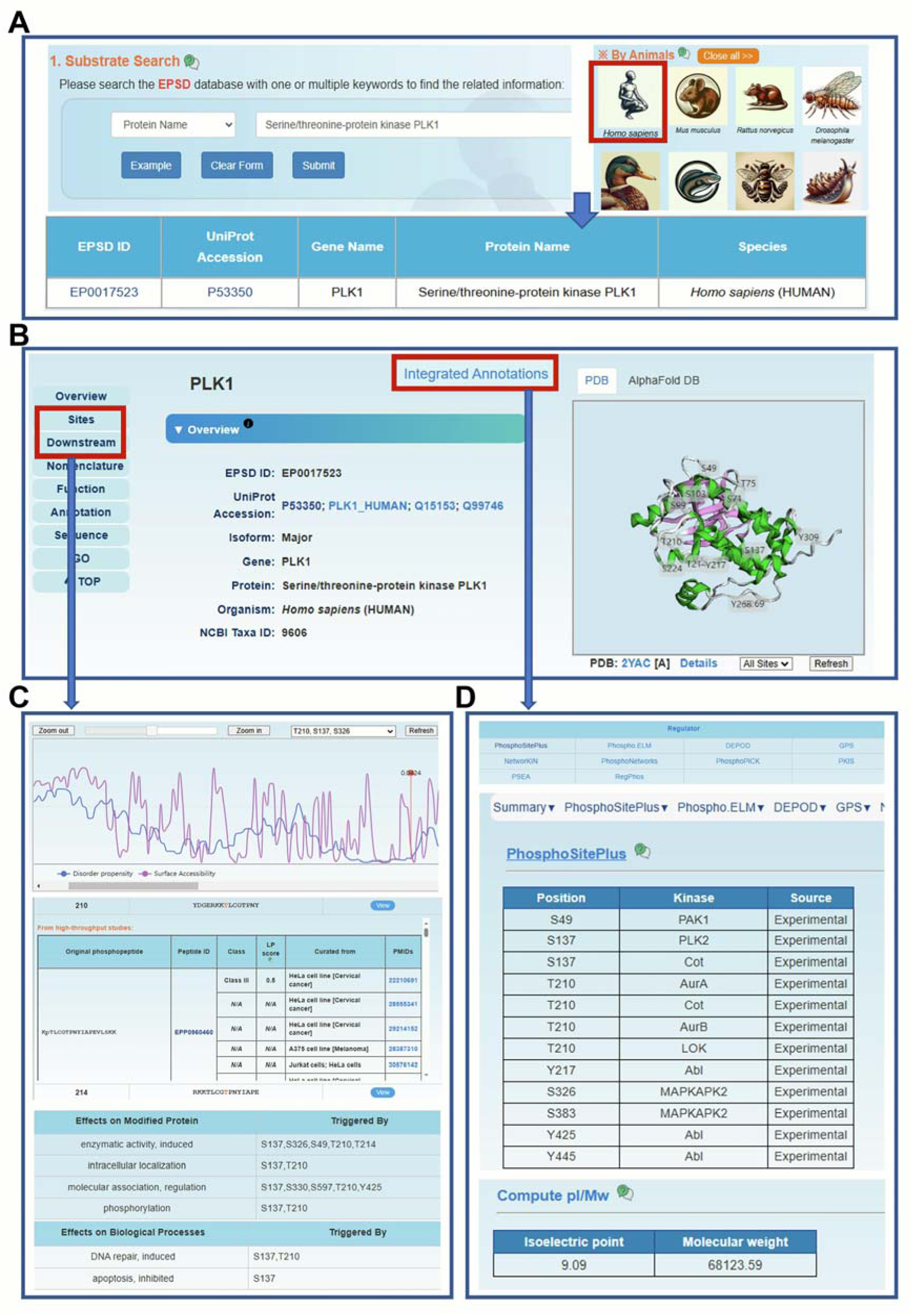
The browse and search option of EPSD 2.0. **A**. The “Substrate Search” and “Browse by Species”. **B**. The overview of human PLK1. **C**. The basic information and details on p-sites. **D**. The annotation of PLK1. “Regulator” was presented as an example.

The “View” page typically comprises ten sections. In the “Overview” section, we offer the basic information, including UniProt ID, protein/gene name, organism, and NCBI taxa ID. In addition, if the protein was mapped to the PDB database [56] or AlphaFold database [54], its typical 3D structure would be shown by 3Dmol.js [57] with p-sites highlighted on the structure (Figure 3B). In the section of “Sites”, users can view the p-sites with corresponding disorder propensity and surface accessibility calculated by IUPred [58] and NetSurfP[59], respectively. Previous research has shown that p-sites are predominantly situated in intrinsically disordered regions [60,61], and the residues with higher surface accessibility are more likely to be phosphorylated [62,63]. In turn, phosphorylation can result in conformational changes of proteins, which may alter disorder regions and surface accessibility [61,64]. Therefore, we consider this information will assist biologists in identifying and prioritizing p-sites with potential functional significance. Additionally, all the p-sites are presented in a tabular list along with corresponding source information in the next “Experimentally identified p-sites” section. The reference information is categorized into three types: HTP, Database, and LTP. A “View” button allows users to display further details such as position, phosphopeptides, LP scores, cell/tissue sources, data sources and PMIDs (Figure 3C). For p-sites with information on downstream effects, users can view these annotations along with PMIDs in the section of “Downstream” (Figure 3C). The remaining sections provide annotations for functional descriptions, protein and gene sequences, GO terms and keywords.

Users can access further detailed annotations for this protein in the “Annotation” section or by clicking the “Integrated Annotations” button at the top of the “Overview” section (Figure 3B). Subsequently, clicking on each category will redirect to the page that displays the relevant annotation content. For instance, users can click on “PhosphoSitePlus” under the “Regulator” title to access protein kinases that phosphorylate protein PLK1 curated from PhosphoSitePlus database [24] (Figure 3D). On the page of annotation, various types of annotation can be accessed through the left column (Figure 3D). For example, the physicochemical property can also be viewed by clicking the option “Physicochemical” on the left (Figure 3D). All the detailed information for the 15 aspects of annotation can be viewed in Figure 4, using human PLK1 as an example.

**Figure 4.**
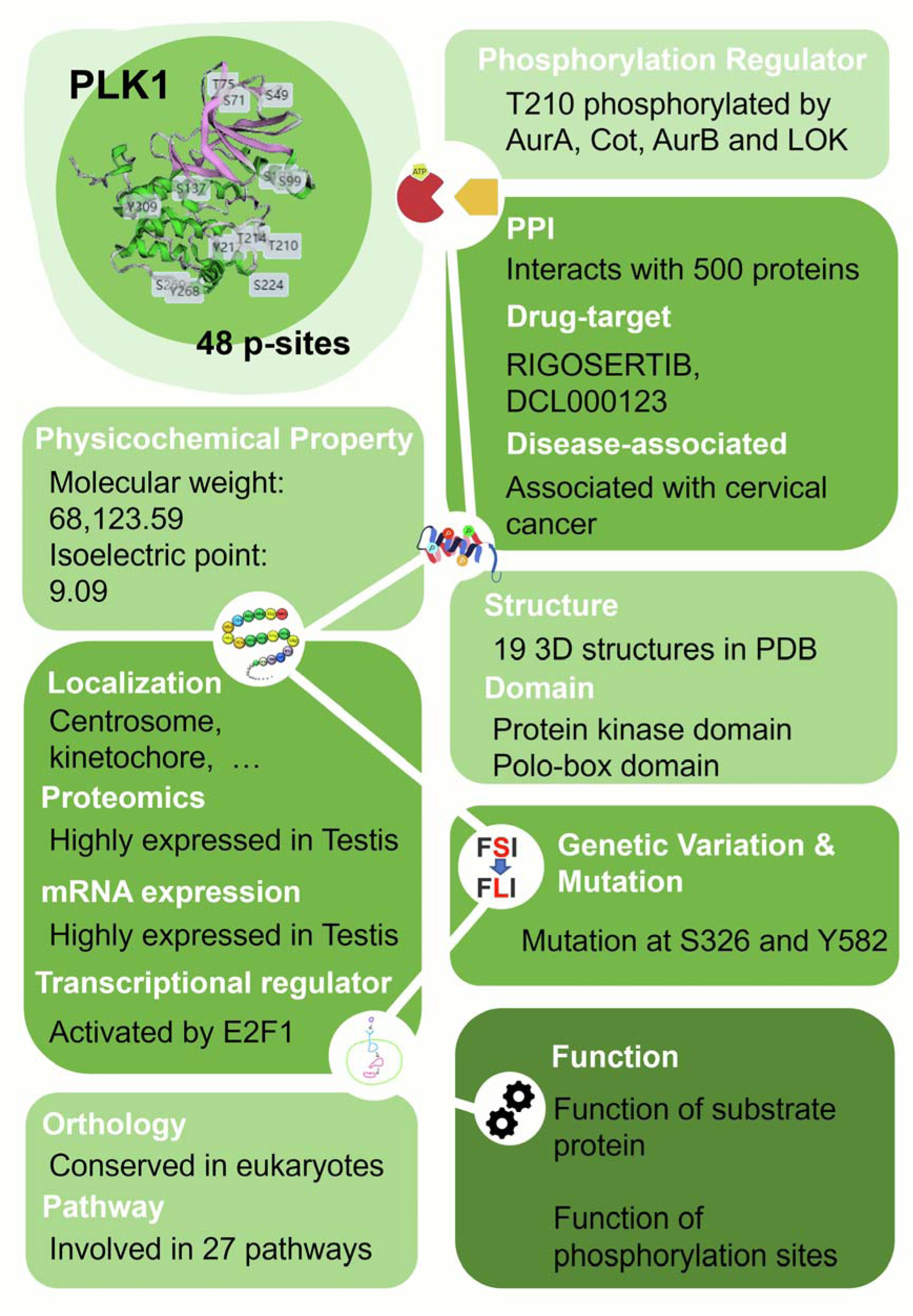
The overview example of integrated annotations. The overview of integrated annotations for human PLK1 protein. A brief summary of all the data resources used in this study is shown in Supplementary Table S2. The details about the processing of each resource were shown in the Supplementary methods.

In addition to the protein search, the search page also offers four other options, such as “Advanced Search”, “Peptide Search”, “Batch Search”, and “BLAST Search”. Each option is tailored to provide different functionalities for more convenient searching. The “Peptide Search” allows users to search the curated phosphopeptides such as “KKpTLCGTPNYIAPEVLSK”, where the “p” indicates that the following amino acid is a p-site. The “Advanced Search” allows users to enter multiple conditions to perform more precise database queries. The “Batch Search” option permits multiple input keywords in a line-by-line format. Moreover, the “BLAST Search” option enables the submission of a single protein sequence in FASTA format to identify identical or homologous proteins in the database.

## Discussion

Protein phosphorylation serves as a crucial and well-studied PTM that modulates protein function, activity, localization, stability, and interactions with other molecules, thereby playing an essential role in virtually all cellular processes [2–7,63–66]. In recent years, MS/MS has continuously generated a number of p-sites across various species [12,13]. The collection and integration of these p-sites can offer valuable resources for elucidating phosphorylation’s regulatory mechanisms. Previously, we developed EPSD 1.0 to curate and annotate p-sites. As the biological research delves deeper into more specific mechanism of biological questions, the functional annotation of p-sites has become increasingly important. In the latest version, EPSD 2.0 introduced significant improvements, particularly in the augmentation of p-sites and inclusion of their functional annotations. Totally, EPSD 2.0 contained 2,769,163 p-sites occurring on 362,707 unique proteins across 223 species, and annotated 88,074 downstream functional events of 32,762 p-sites, covering 58 types of downstream effects on phosphoproteins and 107 regulatory impacts on biological processes. With a total size of 36.2 GB, EPSD 2.0 shows a 2.5-fold expansion in data volume compared to EPSD 1.0 (Supplementary Table S5).

The information in EPSD 2.0 also reveals some interesting insights about phosphorylation. For example, analyzing the downstream effects of p-sites on human PLK1 reveals that the phosphorylation of T210 can trigger lots of effects, such as inducing its own enzymatic activity, altering intracellular localization, and inducing autophagy. Surprisingly, T210 phosphorylation can both induce and inhibit carcinogenesis, which may seem counterintuitive at first glance. By reviewing the original literature, this paradox can be explained. PLK1 is often regarded as a key oncogenic protein because of its critical function in promoting tumor cell division [67]. For example, Zhu et al. discovered that VRK2 kinase phosphorylated PLK1 at T210 to prevent its ubiquitin-dependent proteasomal degradation [68]. This led to the overexpression of PLK1, which drives cancer progression [69]. However, some scientists found that PLK1 overexpression could induce chromosomal instability and suppress tumor development [70]. Yu et al. demonstrated that PLK1 phosphorylated at T210 could act as a tumor suppressor by maintaining the integrity of the spindle assembly checkpoint, preventing chromosomal instability, and inhibiting tumor development in the presence of functional DAB2IP [71]. Understanding these complex regulatory mechanisms is important for the development of effective anti-cancer strategies and for gaining insight into cell cycle regulation. This type of data is important, but its collection is still insufficient.

Although EPSD 2.0 maintained 2,769,163 p-sites, only 32,762 p-sites (1.18%) have been annotated with downstream effects. Thus, the functional effects of most of the p-sites remain to be studied. For example, the S187 residue of human partitioning defective 3 homolog (PARD3, EPSD ID: EP0065790) has been identified as a p-site from multiple phosphoproteomic studies. This site is evolutionarily conserved, and its orthologous counterpart in *D. melanogaster* is S201 of Bazooka (EPSD ID: EP0024205), which was also annotated as an HTP p-site. Based on these annotations, Loyer et al. further uncovered that CDK1 phosphorylates the S201 site of Bazooka to orchestrate neuroblast polarization and sensory organ formation [72] (Figure S1A). In *A. thaliana*, Fu et al. identified phosphorylation of protein MALE DISCOVERER 1 (MDIS1, EPSD ID: EP0151139) at S377, an HTP p-site annotated in EPSD, regulates the auto-phosphorylation of its co-receptor, MDIS1-interacting receptor like kinase 1 (MIK1) [73] (Figure S1B). In addition, Hansen et al. used all annotated phosphoproteins in *M. musculus* from EPSD, and revealed that phosphorylation changes of placental proteins occurred much earlier than mRNA expression changes in response to pulmonary maternal inflammation [74] (Figure S1C). Besides providing useful clues for further experimental design, EPSD also contributes a helpful dataset toward the development of highly accurate predictors. For example, using the data from EPSD 1.0, we previously developed an online service of Group-based Prediction System (GPS) 6.0, which can hierarchically predict kinase-specific p-sites for up to 44,046 protein kinases in 185 eukaryotic species [75]. We believe that more useful predictors will be developed based on the data of EPSD 2.0.

EPSD 2.0 will be regularly maintained and continuously updated to include newly reported p-sites in the literatures. Additionally, more downstream functional events will be collected after reviewing more articles due to the importance of the downstream effects of p-sites and the shortage of their curation. Also, further annotations from external public resources will be included to enhance the comprehensiveness and connectivity of the database.

## Data availability

EPSD 2.0 is accessible at https://epsd.biocuckoo.cn/.

## CRediT author statement

**Miaomiao Chen:** Methodology, Software, Investigation, Data curation, Visualization, Writing – original draft, Writing – review & editing. **Yujie Gou:** Investigation, Data curation, Visualization, Writing – review & editing. **Ming Lei:** Data curation. **Leming Xiao:** Data curation. **Miaoying Zhao:** Data curation. **Xinhe Huang:** Data curation. **Dan Liu**: Data curation. **Zihao Feng:** Data curation. **Di Peng:** Writing – original draft, Writing – review & editing. **Yu Xue:** Writing – original draft, Writing – review & editing, Supervision, Project administration, Funding acquisition. All authors read and approved the final manuscript.

## Acknowledgments

This work was supported by National Key R & D Program of China (2022YFC2704304, and 2021YFF0702000), Natural Science Foundation of China (32341020, 32341021, and 31930021), Hubei Innovation Group Project (2021CFA005), Interdisciplinary Research Program of HUST (2023JCYJ010 and 2024JCYJ013), Hubei Province Postdoctoral Outstanding Talent Tracking Support Program, and Research Core Facilities for Life Science (HUST).

## Declaration of AI and AI-assisted technologies in the writing process

During the preparation of this work the author(s) used DALL·E in order to generate organism images for the web browsing interface. After using this tool/service, the author(s) reviewed and edited the content as needed and take(s) full responsibility for the content of the publication.

## Supplementary material

**Supplementary Figure S1 Three biological examples of the practical research potential of EPSD 2.0.**

**Supplementary Table S1 A summary of mainstream p-site databases for eukaryotic phosphorylation.**

**Supplementary Table S2 The public data resources that included 10 phosphorylation databases and 100 additional databases or tools.**

**Supplementary Table S3 The distribution of p-sites and phosphoproteins across the 223 species.**

**Supplementary Table S4 The effect types of 88,074 functional events.**

**Supplementary Table S5 The comparison of EPSD 2.0 and EPSD 1.0.**

