## Supplementary Text, Figures, and Tables for "EPSD 2.0: An Updated Database of Protein Phosphorylation Sites across Eukaryotic Species": Supplementary Text.docx

***Running title:*** *Chen M and Gou Y / A database of eukaryotic phosphorylation sites*

**Supplementary Data Index**

**Supplementary methods**	**3**

**Supplementary references**	**40**

**Supplementary Tables**	**46**

**Supplementary methods**

The EPSD 2.0 database was implemented in JavaScript, PHP and MySQL. We tested the online service of EPSD 2.0 on a number of mainstream internet browsers, including Microsoft Edge 127, Mozilla Firefox 61.0.2, Google Chrome 128, Opera 55.0 under Windows 10 Operating Systems and Safari 11.1.2 of Apple Mac OS 10.12 (Sierra).

In addition to the manual collection of experimentally identified phosphorylation sites (p-sites) from high-throughput phosphoproteomic studies, EPSD integrated knowledge from 110 existing resources, including 10 phosphorylation databases and 100 additional databases or computational tools (Supplementary Table S2). We carefully processed the data of each resource, and details are shown below.

**1. Public phosphorylation resources**

*1) PhosphoSitePlus (*[*http://www.phosphosite.org/*](http://www.phosphosite.org/)*) [**1**]*

PhosphoSitePlus is one of the best curated and annotated data resources for protein post-translational modifications (PTMs), including phosphorylation, ubiquitylation, acetylation, sumoylation, succinylation, methylation, glycosylation and caspase cleavage, etc [1]. The file ‘Phosphorylation_site_dataset’ that contained experimentally identified p-sites was downloaded from the ‘Downloads’ page of PhosphoSitePlus (on October 23, 2023) [1]. Phosphopeptides in the column ‘SITE_+/-7_AA’ and the corresponding species information in the column ‘ORGANISM’ were extracted. For each organism, phosphopeptides were mapped to canonical sequences and then to isoform sequences, and we obtained 378,691 non-redundant p-sites in 42,245 phosphoproteins.

*2) iPTMnet* (*https://research.bioinformatics.udel.edu/iptmnet/) [**2**]*

iPTMnet is a protein post-translational modification database for PTM knowledge discovery, employing an integrative bioinformatics approach-combining text mining, data mining, and ontological representation to capture rich PTM information []. The ‘iptmnet_ptm’ file containing known p-sites was downloaded from the ‘Download’ page of iPTMnet (Version 6.2, on Octorber 23, 2023) [2]. Only “PHOSPHORYLATION” rows were retained. For each entry, the positions of the p-sites, the UniProtAC of substrate, and the PMIDs were reserved. In total 550,477 non-redundant p-sites of 70,371 proteins were obtained.

*3) RegPhos (*[*http://140.138.144.141/~RegPhos/*](http://140.138.144.141/~RegPhos/%20) *) [**3**]*

RegPhos is a resource to explore the protein kinase-substrate phosphorylation networks [3]. Here, the p-sites data were downloaded from the ‘Download’ page of RegPhos (Version 2.0, on Octorber 23, 2023) [3], containing 3 files from 3 species, including *H. sapiens*, *M. musculus* and Rattus norvegicus. For each entry, we reserved ‘uniproit id’, ’position’, ‘code’ and ‘reference’. In total, 81,335 non-redundant p-sites of 17,214 proteins were obtained.

*4) BioGRID (*[*https://thebiogrid.org/*](https://thebiogrid.org/)*) [**4**]*

BioGRID is a mainstream database for collecting and integrating protein-protein interactions (PPIs) from both small-scale and high-throughput experiments. It also contains a large number of PTM substrates and sites from several model organisms [4]. The ‘BIOGRID-PTMS-LATEST.ptm.zip’ file was downloaded from the ‘downloads’ page of BioGRID (Version 4.4, on November 02, 2023) [4]. For each entry, the column ‘Position’ was used to find positions of p-sites, whereas phosphopeptides centred on phosphorylated S/T or Y residues flanked by 7 residues upstream and 7 residues downstream were retrieved from the protein sequence in the column ‘Sequence’. The corresponding species information and references were taken from the columns ‘Organism Name’ and ‘Pubmed ID’. These phosphopeptides were mapped to canonical sequences and then to isoform sequences, and we obtained 57,394 non-redundant p-sites of 14,520 proteins.

*5) dbPTM (*[*http://dbptm.mbc.nctu.edu.tw*](http://dbptm.mbc.nctu.edu.tw/index.php)*) [**5**]*

dbPTM is a comprehensive database that covers > 130 types of PTMs, and also curated PTM-disease associations, as well as PTM crosstalk events [5]. The compressed file ‘Phosphorylation.zip’ was downloaded from the ‘DOWNLOAD’ page of dbPTM (on Octorber 23, 2023) [5]. Phosphopeptides in the last column and the corresponding species information in the first column were extracted, while the corresponding references were retrieved from the fifth column. For each organism, phosphopeptides were mapped to canonical sequences and then to isoform sequences. In total, 1,614,711 non-redundant p-sites of 226,982 proteins were obtained.

*6) Plant PTM Viewer (https://www.psb.ugent.be/webtools/ptm-viewer/) [**6**]*

Plant PTM Viewer is centralized resource for plant post-translational modifications (PTMs) intuitive for wet- and dry-lab scientists [6]. The file ‘ph_allSPECIES.csv’ that contained experimentally identified p-sites was downloaded from the ‘https://www.psb.ugent.be/webtools/ptm-viewer/download.php’ of Plant PTM Viewer web (on October 23, 2023) [6]. Phosphopeptides in the column ‘PSP’ and the corresponding species information in the column ‘species’ were extracted. For each organism, protein_id was mapped to the corresponding UniProt ID, and we obtained 266,086 non-redundant p-sites in 82,207 phosphoproteins.

*7) Pf-phospho (http://202.54.249.134/DB/index.php/) [**7**]*

Pf-phospho is a resource available for ML-based prediction of phospho-signaling networks of Plasmodium, and it have been benchmarked on the 3098 mass-spectrometry derived phosphorylation sites of Plasmodium species [7]. The file ‘Datasets.xlsx’ that contained experimentally identified p-sites was downloaded from the ‘Database’ page of Pf-phospho (on October 24, 2023) [7]. The protein ids were mapped to the UniProt database, and finally obtained 3,386 non-redundant p-sites in 1,250 phosphoproteins.

*8) Scop3P (https://iomics.ugent.be/scop3p) [**8**]*

Scop3P presents a unique resource for visualization and analysis of phosphosites, and for understanding of phosphosite structure-function relationships. [8]. All the relative files were downloaded from the ‘https://iomics.ugent.be/scop3p/documentation’ page of Scop3P (on October 25, 2023) [8]. ‘Unip_ID’ in file ‘Modification.txt’ was mapped to the file ‘Protein.txt’ to get the corresponding ‘Unip_acc’. We obtained 93,690 non-redundant p-sites in 13,637 phosphoproteins.

*9) PhosPhAt (*[*http://phosphat.uni-hohenheim.de/*](http://phosphat.uni-hohenheim.de/)*) [**9**]*

The Arabidopsis Protein Phosphorylation Site Database (PhosPhAt) integrates and annotates experimentally characterized *Arabidopsis* p-sites [10]. The file ‘Phosphat_20221017.csv’ was downloaded from the ‘Bulk Downloads’ page of PhosPhAt (Version 4.0, on Octorber 23, 2023) [9]. Phosphopeptides of *A. thaliana* in the column ‘modifiedsequence’ were extracted, and the corresponding references were extracted from the column ‘PubMed’. The phosphopeptides were mapped to canonical sequences and then to isoform sequences, and we obtained 49,667 non-redundant p-sites of 11,620 plant proteins.

*10) UniProt (*[*http://www.uniprot.org/*](http://www.uniprot.org/)*) [**11**]*

The compressed file ‘uniprot_sprot.dat’ was downloaded from the FTP server of UniProt ‘ftp://ftp.uniprot.org/pub/databases/uniprot/current_release/knowledgebase/complete/’ (on Octorber 31, 2023) [11]. Phosphothreonine, phosphothreonine and phosphotyrosine residues were retrieved from the lines that started with ‘FT MOD_RES’ for each protein entry if available. To ensure the data quality, p-sites annotated with ‘By similarity’, ‘Potential’ or ‘Probable’ were excluded. Then, for each phosphoprotein, phosphopeptides centred on pS/pT/pY residues flanked by 7 residues upstream and 7 residues downstream were retrieved based on the protein sequences and positions of p-sites, while the corresponding references were also extracted for known p-sites. In total, we obtained 66,490 non-redundant p-sites of 18,466 proteins.

**2. Phosphorylation regulator**

*14) PhosphoSitePlus (*[*http://www.phosphosite.org/*](http://www.phosphosite.org/)*) [**1**]*

The compressed file ‘Kinase_Substrate_Dataset’ was downloaded from the ‘Downloads’ page of PhosphoSitePlus (on Octorber 23, 2023) [1]. Known kinase-substrate relations across different organisms were filtered out directly. The gene names of protein kinases and UniProt IDs of substrates were extracted from the columns of ‘KINASE’ and ‘SUB_ACC_ID’, whereas phosphopeptides in substrates were taken from the column ‘SITE_+/-7_AA’. The UniProt IDs were used to find phosphorylated substrates in the EPSD, while phosphopeptides were mapped to their corresponding protein sequences to pinpoint the positions of the p-sites.

*15) Phospho.ELM (*[*http://phospho.elm.eu.org/*](http://phospho.elm.eu.org/)*) [**12**]*

The file ‘phosphoELM’ was downloaded from the ‘Download’ page of Phospho.ELM (Version 9.0, on Octorber 23, 2023) [12]. The gene names of protein kinases and UniProt IDs of substrates were extracted from the columns of ‘kinases’ and ‘acc’. For each entry, phosphopeptides centred on pS/pT/pY residues flanked by 7 residues upstream and 7 residues downstream were generated from the protein sequence in the column ‘sequence’, as well as the information of p-site positions in the column ‘position’ Then, UniProt IDs were used to find phosphorylated substrates in the EPSD, while phosphopeptides were mapped to their corresponding protein sequences to pinpoint the positions of p-sites.

*16) PostMod (*[*http://pbil.kaist.ac.kr/PostMod/*](http://pbil.kaist.ac.kr/PostMod/)*) [**13**]*

The training dataset of PostMod contained in the compressed file ‘uploadDataset’ was downloaded from the ‘Database’ page of PostMod (on March 11, 2024) [13]. For each positive dataset, the names of the protein kinases or kinase groups were extracted from the last column, whereas UniProt IDs listed in the first column were mapped to EPSD.

*17) PSEA (*[*http://bioinfo.ncu.edu.cn/PKPred_Home.aspx*](http://bioinfo.ncu.edu.cn/PKPred_Home.aspx)*) [**14**]*

The compressed file ‘Kinase-specific phosphorylation data’ was downloaded from the ‘DOWNLOAD’ page of PSEA (on March 11, 2024) [14]. The gene names of protein kinases and UniProt IDs of substrates were extracted from the columns of ‘Kinase’ and ‘Substrate ACC’, whereas phosphopeptides in substrates were taken from the column ‘Sequence(-7~+7)’. Then, UniProt IDs were adopted to find phosphorylated substrates in the EPSD, while phosphopeptides were mapped to their corresponding protein sequences to pinpoint the positions of p-sites.

*18) PhosphoNetworks (*[*http://www.phosphonetworks.org/*](http://www.phosphonetworks.org/)*) [**15**]*

The file ‘refKSI’ was downloaded from the ‘Download’ page of PhosphoNetworks (on April 09, 2024) [15]. The gene names of protein kinases and substrates were extracted from the columns of ‘kinase’ and ‘substrate’. In PhosphoNetworks [15], exact phosphorylation sites in substrates were not provided, while only kinase-substrate relations in *H. sapiens* were present. We directly mapped the gene names of substrates to human phosphoproteins in EPSD.

*19) HuPHO (*[*http://hupho.uniroma2.it/*](http://hupho.uniroma2.it/)*) [**16**]*

The file ‘Substrates_2019-05-13_01-58’ was downloaded from the ‘Resources & Tools’ page of HuPHO (on May 13, 2019) [16]. The UniProt IDs and gene names of human phosphatases in the columns of ‘UniprotKB-AC Pho’ and ‘Gene Name HS’ were extracted, respectively. The UniProt IDs of substrates were retrieved from the column ‘UniprotKB-AC Sub’, whereas the corresponding references were extracted from the columns of ‘PMID in vivo’, ‘PMID in vitro’ and ‘PMID trapping’.

*20) DEPOD (*[*http://www.depod.bioss.uni-freiburg.de/*](http://www.depod.bioss.uni-freiburg.de/)*) [**17**]*

The human DEPhOsphorylation Database (DEPOD) is a manually curated resource for human active phosphatases. The ‘PPase_protSubstrates_201903.xllls’ file was downloaded from the ‘Download’ page of DEPOD (on April 9, 2024) [17]. The gene names of phosphatases and substrates in the columns of ‘Phosphatase’ and ‘Substrate’ were extracted, respectively. DEPOD provided dephosphorylation sites for a considerable number of human substrates, and known dephosphorylation sites were obtained from the column ‘Dephosphorylation site’, if available. For each pair of known phosphatase-substrate relation, the corresponding references were extracted from the column ‘PubMed ID’. We mapped gene names of substrates to human phosphoproteins in EPSD.

*21) GPS (*[*http://gps.biocuckoo.org/*](http://gps.biocuckoo.org/)*) [**18**]*

Previously, we developed a software package of Group-based Prediction System (GPS) for the prediction of kinase-specific p-sites from protein sequences [18]. In this work, GPS 3.0 was used to predict potentially regulatory kinases of known p-sites in 8 model organisms, including *H. sapiens*, *M. musculus*, *R. norvegicus*, *D. melanogaster*, *C. elegans*, *A. thaliana*, *S. pombe* and *S. cerevisiae*. For a better coverage, high thresholds of GPS with false positive rates of 2% for serine/threonine kinases and 4% for tyrosine kinases were adopted. For each protein entry, details were provided with p-site positions (‘Position’), p-site types (‘Code’), predicted kinases (‘Kinase’), flanking peptides (‘Peptide’), predicted scores (‘Score’) and pre-defined threshold values for different kinase groups or families (‘Cutoff’).

*22) NetworKIN (*[*http://www.networkin.info/*](http://www.networkin.info/)*) [**19**]*

The compressed file ‘networkin_human_predictions_3.1.tsv’ was downloaded from the ‘Download’ page of NetworKIN (on May 6, 2019) [19]. This file contained pre-calculated site-specific kinases- or SH2-substrate relations for human known p-sites from the KinomeXplorer database [19], using Ensembl protein sequences (Version 59) [20]. We mapped Ensembl protein accession numbers in the column ‘#substrate’ to human phosphoproteins in EPSD, while phosphopeptides in the column ‘sequence’ were mapped to their corresponding protein sequences to pinpoint the positions of p-sites. Information in columns ‘networkin_score’, ‘tree’, ‘netphorest_group’, ‘netphorest_score’, ‘string_identifier’, ‘string_score’, ‘substrate_name’ and ‘string_path’ were also integrated.

*23) PKIS (*[*http://bioinformatics.ustc.edu.cn/pkis/*](http://bioinformatics.ustc.edu.cn/pkis/)*) [**21**]*

Pre-calculated human kinase-specific p-sites of different kinases were downloaded from PKIS (on May 7, 2019) [21]. The Ensembl protein IDs and p-site positions were extracted from lines started with ‘>’, and mapped to human proteins in EPSD.

*24) PhosphoPICK (*[*http://bioinf.scmb.uq.edu.au/phosphopick/phosphopick*](http://bioinf.scmb.uq.edu.au/phosphopick/phosphopick)*) [**22**]*

The pre-predicted results of potential kinase-substrate relations for *H. sapiens*, *M. musculus* and *S. cerevisiae* were downloaded from the ‘Download’ page of PhosphoPICK (on May 6, 2019) [22]. The UniProt IDs of substrates were extracted from the column ‘protein-accession’, while protein kinase names were taken from the first line. For each substrate, only predicted kinases with *p*-value < 0.05 were kept. PhosphoPICK was designed for the prediction of kinase-substrate relations, not in a site-specific manner [22].

*25) RegPhos (*[*http://140.138.144.141/~RegPhos/index.php*](http://140.138.144.141/~RegPhos/index.php)*) [**23**]*

The kinome annotation files ‘RegPhos_Phos_human.txt’, ‘RegPhos_Phos_mouse.txt’ and ‘RegPhos_Phos_rat.txt’ were downloaded from the ‘Download’ page of RegPhos (on Octorber 23, 2023). The gene names of protein kinases and UniProt IDs of substrates were extracted from the columns of ‘catalytic kinase’ and ‘AC’, while the information of p-sites were generated from the columns of ‘positon’ and ‘code’.

**3. Genetic variation & mutation**

*26) TCGA (*[*https://www.cancer.gov/tcga*](https://www.cancer.gov/tcga)*) [**24**]*

We downlowaded TCGA mutations provided by BROAD Institute ([http://gdac.broadinstitute.org/runs/stddata__latest/data](http://gdac.broadinstitute.org/runs/stddata__2016_01_28/data), Oncotated calls, level 3, in April 2023) [24]. All 36 available projects were downloaded, including adrenocortical carcinoma (ACC), bladder urothelial carcinoma (BLCA), Breast invasive carcinoma (BRCA), cervical squamous cell carcinoma and endocervical adenocarcinoma (CESC), cholangiocarcinoma (CHOL), colon adenocarcinoma (COAD), colorectal cancer (COADREAD), lymphoid neoplasm diffuse large B-cell lymphoma (DLBC), esophageal carcinoma (ESCA), glioblastoma multiforme (GBM), glioma (GBMLGG), head and neck squamous cell carcinoma (HNSC), kidney chromophobe (KICH), pan-kidney cohort (KIPAN, KICH+KIRC+KIRP), kidney renal clear cell carcinoma (KIRC), kidney renal papillary cell carcinoma (KIRP), acute myeloid leukemia (LAML), brain lower grade glioma (LGG), liver hepatocellular carcinoma (LIHC), lung adenocarcinoma (LUAD), lung squamous cell carcinoma (LUSC), ovarian serous cystadenocarcinoma (OV), pancreatic adenocarcinoma (PAAD), pheochromocytoma and paraganglioma (PCPG), prostate adenocarcinoma (PRAD), rectum adenocarcinoma (READ), sarcoma (SARC), skin cutaneous melanoma (SKCM), stomach adenocarcinoma (STAD), stomach and esophageal carcinoma (STES), testicular germ cell tumors (TGCT), thyroid carcinoma (THCA), thymoma (THYM), uterine corpus endometrial carcinoma (UCEC), uterine carcinosarcoma (UCS), and uveal melanoma (UVM). Ensembl transcript IDs were used to map the data from TCGA to EPSD (columns entitled ‘Annotation_Transcript’ in TCGA files). Then, columns including ‘PRJ_code’, ‘Chromosome’, ‘Start_position’, ‘End_position’, ‘Strand’, ‘Variant_Classification’, ‘Variant_Type’, ‘Reference_Allele’, ‘Tumor_Seq_Allele1’, ‘Tumor_Seq_Allele2’, ‘Tumor_Sample_Barcode’, ‘Matched_Norm_Sample_Barcode’, ‘Genome_Change’, ‘cDNA_Change’, ‘Codon_Change’ and ‘Protein_Change’ were integrated. Positions of human cancer mutations in the column ‘Protein_Change’ were used, and only cancer mutations directly changed known p-sites were reserved.

*27) ICGC (*[*http://icgc.org/*](http://icgc.org/)*) [**25**]*

We downloaded all simple somatic mutations of tumour tissues from ICGC data portal (<https://dcc.icgc.org/releases/release_28/Projects/>, release 28, in May 2024) [25]. Ensembl gene IDs (columns entitled ‘gene_affected’ in ICGC data) were used to map mutations to EPSD. Columns including ‘mutation’, ‘project’, ‘chromosome’, ‘start’, ‘end’, ‘mutation_type’, ‘consequence_type’, ‘aa_mutation’ and ‘cds_mutation’ were integrated. Positions of cancer mutations in the column ‘aa_mutation’ were used, while only cancer mutations directly changed known p-sites were reserved. Here, we obtained 182,351 cancer mutations for 13,865 substrate genes.

*28) COSMIC (*[*https://cancer.sanger.ac.uk/cosmic*](https://cancer.sanger.ac.uk/cosmic)*) [**26**]*

We obtained cancer mutations by downloading the file ‘CosmicMutantExportCensus.tsv.gz’ from COSMIC (<https://cancer.sanger.ac.uk/cosmic/download>, release v90, on March, 2020) [26]. Ensembl transcript IDs were used as primary accession numbers for mapping cancer mutations to EPSD. Columns including ‘Sample’, ‘Mutation’, ‘Primary site’, ‘Primary histology’, ‘Mutation CDS’, ‘Mutation AA’, ‘Description’, ‘Position’ and ‘Strand’ were integrated. Positions of mutations in the column ‘Mutation AA’ were used, while only cancer mutations directly changed known p-sites were reserved. Here, we obtained 151,128 cancer mutations for 14,715 substrate genes.

*29) dbSNP (*[*https://www.ncbi.nlm.nih.gov/snp/*](https://www.ncbi.nlm.nih.gov/snp/)*) [**27**]*

25 reference SNP files such as ‘refsnp-chr1.json.bz2’ of *H. sapiens* were downloaded from the ftp site of dbSNP (https://ftp.ncbi.nih.gov/snp/latest_release/JSON/, release November 16, 2022) (28). SNPs identified by ‘refsnp_id’, together with their positions in protein sequences as column ‘AA Position’, corresponding amino acids residues as column ‘AA residue’, variation type as column ‘Type’, positions in nucleotide sequence as column ‘Position’, alleles as column ‘Alleles’ and variation frequency as column ‘Frequency’ were extracted. Positions of nsSNPs in the column ‘AA position’ were used, while only nsSNPs directly changed known p-sites were reserved.

*30) IntOGen (*[*https://www.intogen.org/*](https://www.intogen.org/)*) [**28**]*

We downloaded the catalog of driver mutations from the ‘Downloads’ page of IntOGen (released in May 2023) [28]. Ensembl transcript IDs were used as primary accession numbers for mapping cancer mutations to EPSD. Columns including ‘cancer’, ‘consequence’, ‘gdna’, ‘cdna’ and ‘protein’ were integrated. Positions of mutations in the column ‘protein’ were used, while only cancer mutations directly changed known p-sites were reserved.

*31) MIMP (*[*http://mimp.baderlab.org/*](http://mimp.baderlab.org/)*) [**29**]*

MIMP is a web server to investigate the impact of mutations on kinase-substrate phosphorylation [29]. The file ‘tcga_rewiring_events_prob.tab’ was downloaded from the ‘Download’ page of MIMP (on May 13, 2019). The human gene names were extracted from the column ‘gene’, while original residues, mutant residues and positions of single amino acid mutations (SAAMs) from the columns ‘mut’. Positions of the phosphorylation sites were extracted from the column ‘position’, information in columns ‘flank_wt’, ‘flank_mt’, ‘wt_score’, ‘mt_score’, ‘pwm’, ‘effect’ and ‘log_ratio’ were also considered. Then p-sites were mapped to known p-sites in EPSD according to the gene names and positions.

**4. Functional annotation**

*32) iEKPD (*[*http://iekpd.biocuckoo.org/*](http://iekpd.biocuckoo.org/)*) [**30**]*

Recently, we developed a hierarchical database of iEKPD 2.0, which contained 109,912 protein kinases (PKs), 23,294 protein phosphatases (PPs) and 68,748 proteins containing phosphoprotein-binding domains (PPBDs) in 164 eukaryotic species, including 74 animals, 47 plants and 43 fungi [30]. In this work, PKs, PPs and PPBDs in 8 model organisms, including *H. sapiens*, *M. musculus*, *R. norvegicus*, *D. melanogaster*, *C. elegans*, *A. thaliana*, *S. pombe* and *S. cerevisiae*, were mapped to EPSD by using UniProt IDs.

*33) iUUCD (*[*http://iuucd.biocuckoo.org/*](http://iuucd.biocuckoo.org/)*) [**31**]*

Previously, we developed an updated database iUUCD 2.0, which contained 1,230 ubiquitinactivating enzymes (E1s), 5,636 ubiquitin-conjugating enzymes (E2s), 93,343 ubiquitin-protein ligases (E3s), 9,548 deubiquitinating enzymes (DUBs), 30,173 ubiquitin/ubiquitin-like binding domains (UBDs) and 11,099 ubiquitin-like domains (ULDs) [31]. In this work, ubiquitin and ubiquitin-like regulators in 8 model organisms, including *H. sapiens*, *M. musculus*, *R. norvegicus*, *D. melanogaster*, *C. elegans*, *A. thaliana*, *S. pombe* and *S. cerevisiae*, were mapped to EPSD by using gene names.

*34) WERAM (*[*http://weram.biocuckoo.org/*](http://weram.biocuckoo.org/)*) [**32**]*

In 2017, we developed a database WERAM, which contained histone acetyltransferases (HATs), histone deacetylases (HDACs), histone methyltransferases (HMTs), histone demethylases (HDMs) and acetyl- or methyl-binding proteins in 148 eukaryotes [32]. In this work, histone acetylation and methylation regulators in 8 model organisms, including *H. sapiens*, *M. musculus*, *R. norvegicus*, *D. melanogaster*, *C. elegans*, *A. thaliana*, *S. pombe* and *S. cerevisiae*, were mapped to EPSD by using gene names.

*35) AnimalTFDB (*[*http://bioinfo.life.hust.edu.cn/AnimalTFDB/*](http://bioinfo.life.hust.edu.cn/AnimalTFDB/)*) [**33**]*

The transcription factors (TFs) and transcription cofactors (TF cofactors) were downloaded from the ‘Download’ page of AnimalTFDB (<http://bioinfo.life.hust.edu.cn/AnimalTFDB/#!/download>, Version 4.0, on January, 2023) [33]. In this work, TFs and TF cofactors in 5 model organisms, including *H. sapiens*, *M. musculus*, *R. norvegicus*, *D. melanogaster* and *C. elegans*, were mapped to EPSD with Ensembl gene IDs.

*36) PlantTFDB (*[*http://planttfdb.cbi.pku.edu.cn/*](http://planttfdb.cbi.pku.edu.cn/)*) [**34**]*

We downloaded the TF list of *A. thaliana* from the ‘Download’ page of PlantTFDB (https://planttfdb.gao-lab.org/download.php, Version 5.0, in Dec 2022) [34]. The TF families in the column ‘Family’ were extracted, and Ensembl gene IDs in the column ‘Gene_ID’ were retrieved and mapped to EPSD.

*37) AmyPro (*[*http://amypro.net/*](http://amypro.net/)*) [**35**]*

The amyloid fibre-forming proteins were obtained from the file ‘amypro.fasta‘ on ‘Download’ page of AmyPro (in May 2024) [35]. Columns including ‘ID’, ‘category’, ‘pubmed’ and ‘uniprot’ were integrated, and the UniProt IDs were mapped to EPSD.

*38) HAMAP (*[*http://hamap.expasy.org/*](http://hamap.expasy.org/)*) [**36**]*

High-quality Automated and Manual Annotation of Proteins (HAMAP) maintains a collection of manually curated family profiles for protein classification and annotation [36]. In this work, the file ‘rules_index.dat’ and the compressed file ‘hamap_alignments’ were downloaded from the FTP server (<ftp://ftp.expasy.org/databases/hamap/>) of HAMAP (release 2022_10, in May 2022). The HAMAP rule accession numbers, names and descriptions were extracted from the file ‘rules_index.dat’, whereas. UniProt IDs and rule accession numbers were extracted from the file ‘hamap_alignments’. Then, protein entries were mapped to phosphoproteins in EPSD using UniProt IDs.

*39) Membranome (*[*http://membranome.org/*](http://membranome.org/)*) [**37**]*

The file ‘proteins-2024-05-07.csv’ was downloaded in the ‘DOWNLOAD FILES’ page of Membranome (Version 3.0, in May 2024). The Membranome IDs in the column ‘id’ were extracted, and UniProt IDs in the column ‘uniprotcode’ were retrieved and mapped to EPSD.

*40) neXtProt (*[*https://www.nextprot.org/*](https://www.nextprot.org/)*) [**38**]*

The file ‘nextprot_ac_list_all.txt’ was downloaded from the FTP server (<ftp://ftp.nextprot.org/pub/current_release/ac_lists/>) of neXtProt (Version v2.56.0, in May 2024). For each entry, the neXtProt IDs were extracted, while UniProt IDs were retrieved and mapped to EPSD.

*41) iPCD (https://ipcd.biocuckoo.cn/) [**39**]*

Recently, we developed a comprehensive database of integrated annotations for Programmed Cell Death (iPCD), containing 1,094,627 known and computationally predicted regulators for 31 forms of PCDs in 562 eukaryotes [39]. In this work, iPCD proteins in 8 model organisms, including *H. sapiens*, *M. musculus*, *R. norvegicus*, *D. melanogaster*, *C. elegans*, *A. thaliana*, *S. pombe* and *S. cerevisiae*, were mapped to EPSD by using gene names.

*42) CGDB (*[*http://cgdb.biocuckoo.org/*](http://cgdb.biocuckoo.org/)*) [**40**]*

Previously, we developed a database of circadian genes in eukaryotes, containing ~73,000 circadian-related genes in 68 animals, 39 plants and 41 fungi [40]. In this work, CGDB genes in 8 model organisms, including *H. sapiens*, *M. musculus*, *R. norvegicus*, *D. melanogaster*, *C. elegans*, *A. thaliana*, *S. pombe* and *S. cerevisiae*, were mapped to EPSD by using gene names.

*43) MultitaskProtDB-II (*[*http://wallace.uab.es/multitaskII/*](http://wallace.uab.es/multitaskII/)*) [**41**]*

MultitaskProtDB-II is a repository of multitasking (moonlighting) proteins curated in the literature [41]. The multitasking proteins were exported from the ‘DataBase’ page of MultitaskProtDB-II (<http://wallace.uab.es/multitaskII/proteins_list.php>, in June 2024). Columns including ‘ID’, ‘Uni Prot’, ‘Protein Name’, ‘Canonical Function’, ‘Moonlighting Function’ and ‘Reference’ were integrated, and the UniProt IDs were mapped to EPSD.

*44) MoonDB (*[*http://moondb.hb.univ-amu.fr/*](http://moondb.hb.univ-amu.fr/)*) [**42**]*

The file ‘all_EMF_annotation.tsv’ was downloaded from the ‘Downloads’ page of MoonDB (<http://moondb.hb.univ-amu.fr/downloads>, Version 2.0, in May 2024) [42]. Columns including ‘UniprotKB_AC’, ‘GO_ID_1’, ‘GO_Name_1’, ‘GO_ID_2’, ‘GO_ID_Name’, ‘Association_Prob_E_Value’ and ‘Interaction_Prob_E_Value’ were extracted, and the UniProt IDs were mapped to EPSD. Here, we taken 280 multitasking proteins that can be phosphorylated in 5 model organisms, including *H. sapiens*, *M. musculus*, *D. melanogaster*, *C. elegans* and *S. cerevisiae*.

*45) CORUM (*[*http://mips.helmholtz-muenchen.de/corum/*](http://mips.helmholtz-muenchen.de/corum/)*) [**43**]*

The CORUM database provides a manually curated repository of experimentally characterized protein complexes from mammalian organisms [43]. The file ‘allComplexes.txt.zip’ were obtained from CORUM (<http://mips.helmholtz-muenchen.de/corum/#download>, in May 2024). The UniProt IDs in the column ‘subunits(UniProt IDs)’ were extracted and mapped to EPSD.

*46) CellMarker (*[*http://biocc.hrbmu.edu.cn/CellMarker/*](http://biocc.hrbmu.edu.cn/CellMarker/)*) [**44**]*

The CellMarker database is a manually curated resource of cell markers for various cell types in tissues of human and mouse [44]. The file ‘all_cell_markers.txt’ was downloaded from the ‘Download’ page of CellMarker (http://117.50.127.228/CellMarker/CellMarker_download.html, in June, 2024). Columns including ‘tissueType’, ‘cancerType’, ‘cellType’, ‘cellName’, ‘cellMarker’, ‘markerResource’ and ‘PMID’ were integrated, and the UniProt IDs in the column ‘proteinID’ were extracted and mapped to EPSD.

*47) GPCRdb (*[*http://www.gpcrdb.org/*](http://www.gpcrdb.org/)*) [**45**]*

GPCRdb contains data, diagrams and web tools for G protein-coupled receptors (GPCRs) [45]. The file ‘uniprot_mapping.txt’ was downloaded from the ‘Tutorial’ page of GPCRdb (<http://docs.gpcrdb.org/linking.html>, Release May, 2024). The UniProt IDs were extracted with GPCRdb IDs, and mapped to EPSD.

*48)* *PTMCode (*[*http://ptmcode.embl.de/*](http://ptmcode.embl.de/)*) [**46**]*

PTMcode is a database focusing on the functional associations between protein PTMs [46]. The compressed files ‘PTMcode2_associations_within_proteins.txt’ and ‘PTMcode2_associations_between_proteins.txt’ were downloaded from the ‘Data’ page of PTMCode (on Dec 12, 2022). Gene names and the corresponding species information were extracted from the columns ‘Protein1’, ‘Protein2’ and ‘Species’, while types and positions of PTM sites were retrieved from the columns ‘PTM1’, ‘PTM2’, ‘Residue1’ and ‘Residue2’. The data was mapped to EPSD by using gene names, and PTM functional associations with at least one known p-site were reserved.

**5. Structural annotation**

*49) PDB (*[*http://www.rcsb.org/*](http://www.rcsb.org/pdb/home/home.do)*) [**47**]*

From the FTP server (<ftp://ftp.rcsb.org/pub/pdb/data/structures/divided/pdb/>) of PDB, 150,037 .ent files was downloaded (in June 2024) [47]. For each .ent file, the PDB entry, UniProt ID, protein sequence, title and chain were extracted. Then, the PDB entries were mapped to phosphoproteins according to the UniProt IDs.

*50) MMDB (*[*https://www.ncbi.nlm.nih.gov/structure/*](https://www.ncbi.nlm.nih.gov/structure/)*) [**48**]*

We downloaded the file ‘nrpdb.gz’ from FTP server of MMDB (<ftp://ftp.ncbi.nih.gov/mmdb/nrtable/>, in Oct 2022). From the PDB database [47], we obtained all PDB IDs together with corresponding UniProt IDs. Then using PDB IDs, we mapped the MMDB data to EPSD. The columns ‘PDB code’, ‘Chain ID’, ‘MMDB ID’, ‘Resolution’, ‘Method of coordinate determination’ and ‘Acceptable in structural quality’ were reserved for the integration.

*51) SCOP2 (*[*http://scop2.mrc-lmb.cam.ac.uk/*](http://scop2.mrc-lmb.cam.ac.uk/)*) [**49**]*

We downloaded the files ‘scop2_nodes_names_20140205’, ‘domain_segments_pdb_20140205’, ‘domain_segments_seq_20140205’, and ‘domains2nodes_20140205’ from download page of SCOP2 (<http://scop2.mrc-lmb.cam.ac.uk/downloads/>, in February 2018) [49]. We obtained functional domain IDs and locations on proteins from ‘domain_segments_seq_20140205’. We extracted PDB IDs and exact positions of SCOP domains from ‘domain_segments_pdb_20140205’. SCOP node IDs for all domains were taken from ‘domains2nodes_20140205’. Furthermore, we extracted names for mapped nodes from ‘scop2_nodes_names_20140205’. Finally, the columns ‘Domain’, ‘Domain_serial’, ‘Node’, ‘Name’, ‘PDB’, ‘Chain’, ‘BeginPDB’, ‘EndPDB’, ‘UniprotKB’, ‘BeginUni’ and ‘EndUni’ were retained for integration. The data in SCOP2 was mapped to EPSD by using UniProt IDs in the column ‘UniprotKB’. Here, we obtained 922 entries of 245 phosphoproteins of the 8 model organisms.

*52) IUPred (*[*https://iupred.elte.hu/*](https://iupred.elte.hu/)*) [**50**]*

IUPred was designed for the prediction of disordered regions from protein sequences [50]. Here, the program package was downloaded from the ‘Downloads’ page of IUPred (version 3.0), to predict potentially disordered regions in phosphoproteins of 8 model organisms (on Oct 21, 2022).

*53) MobiDB (*[*http://mobidb.bio.unipd.it/*](http://mobidb.bio.unipd.it/)*) [**51**]*

The MobiDB database provides protein disorder and mobility annotations [51]. The compressed file ‘all_disorder_sprot.mjson’ was downloaded from the ‘Datasets’ page of MobiDB (<http://mobidb.bio.unipd.it/dataset>, Version 5.0, in Oct 2022). The disorder content and regions were integrated, and the UniProt IDs in the column ‘acc’ were extracted and mapped to EPSD.

**6. Physicochemical property**

*54) AAindex (*[*http://www.genome.jp/aaindex/*](http://www.genome.jp/aaindex/)*) [**52**]*

AAindex is a resource that contains various biochemical and physicochemical properties of amino acids. The file ‘aaindex1’ was downloaded from the FTP server (ftp://ftp.genome.jp/pub/db/community/aaindex/, on May 8, 2019). For each physicochemical property, scores for 20 types of amino acids were extracted.

*55) Compute pI/Mw (*[*https://web.expasy.org/compute_pi/*](https://web.expasy.org/compute_pi/)*) [**53**]*

The web service of Compute pI/Mw was adopted for the computation of theoretical isoelectric point (pI) and molecular weight (Mw) for all phosphoproteins of 8 model organisms (on May 21, 2024).

**7. Functional domain**

*56) Pfam (*[*http://pfam.xfam.org/*](http://pfam.xfam.org/)*) [**54**]*

The file ‘‘Pfam-A.regions.uniprot.tsv.gz’ was downloaded from the FTP server (<ftp://ftp.ebi.ac.uk/pub/databases/Pfam>) of Pfam (in May 2024). For each entry, the Pfam and UniProt IDs in the columns ‘Pfam ID’ and ‘uniprot_acc’ were extracted. Then Pfam entries were mapped to phosphoproteins in EPSD according to UniProt IDs.

*57) PROSITE (*[*https://prosite.expasy.org/*](https://prosite.expasy.org/)*) [**55**]*

The compressed file ‘prosite_alignments.tar.gz’ was downloaded from the FTP server (<ftp://ftp.expasy.org/databases/prosite/>) of PROSITE (on May, 2024). For each entry, the PROSITE ID, UniProt IDs and corresponding descriptions were extracted. Then PROSITE entries were mapped to phosphoproteins according to UniProt IDs.

*58) InterPro (*[*http://www.ebi.ac.uk/interpro/*](http://www.ebi.ac.uk/interpro/)*) [**56**]*

The compressed file ‘protein2ipr.dat’ was downloaded in the ‘Download’ page of InterPro (<http://www.ebi.ac.uk/interpro/download.html>, Version 2023_05, in June 2024). For each entry, the InterPro ID, UniProt IDs, family name, start and end positions were retrieved. Then InterPro entries were mapped to phosphoproteins according to UniProt IDs.

*59) Gene3D (*[*http://gene3d.biochem.ucl.ac.uk/*](http://gene3d.biochem.ucl.ac.uk/)*) [**57**]*

From the FTP server (<ftp://orengoftp.biochem.ucl.ac.uk/gene3d/CURRENT_RELEASE/>) of Gene3D (Version 16.0, in April 2019), the compressed files ‘model_to_family_map.csv’ and ‘representative_uniprot_genome_assignments_with_eval.csv’ were downloaded. The Gene3D IDs and descriptions were extracted from the file ‘model_to_family_map.csv’, whereas corresponding UniProt IDs were retrieved from the file ‘representative_uniprot_genome_assignments_with_eval.csv’. Then, Gene3D entries were mapped to phosphoproteins according to UniProt IDs.

*60) PIRSF (*[*https://proteininformationresource.org/pirwww/dbinfo/pirsf.shtml*](https://proteininformationresource.org/pirwww/dbinfo/pirsf.shtml)*) [**58**]*

The ‘pirsfinfo.dat’ file was downloaded from the FTP server (<ftp://ftp.pir.georgetown.edu/databases/pirsf/>) of PIRSF (in June 2024). For each entry, the ‘PIRSF Number’, UniProt IDs and ‘PIRSF Name’ were exacted. Then, PIRSF entries were mapped to phosphoproteins according to UniProt IDs.

*61) PRINTS (*[*http://130.88.97.239/PRINTS/index.php*](http://130.88.97.239/PRINTS/index.php)*) [**59**]*

The compressed file ‘prints42_0.dat’ was downloaded from the FTP server (<ftp://ftp.ebi.ac.uk/pub/databases/prints/>) of PRINTS (Version 42.0, in June 2024). The PRINTS IDs, UniProt IDs, identifiers and titled were extracted. Then, PRINTS entries were mapped to phosphoproteins according to UniProt IDs.

*62) SMART (*[*http://smart.embl-heidelberg.de/*](http://smart.embl-heidelberg.de/)*) [**60**]*

Online SMART batch retrieval tool (<http://smart.embl-heidelberg.de/smart/batch.pl>) was used to obtain the domain information including domain name, start and end positions, and E-value for each phosphoprotein in 8 model organisms, by using UniProt IDs (in June 2024).

**8. Disease-associated information**

*63) ClinVar (*[*https://www.ncbi.nlm.nih.gov/clinvar/*](https://www.ncbi.nlm.nih.gov/clinvar/)*) [**61**]*

First, we downloaded the file ‘clinvar_20240514.vcf’ from the FTP server of ClinVar (<ftp://ftp.ncbi.nlm.nih.gov/pub/clinvar/>, on May 17, 2024). Because ClinVar only provided genomic positions of SNPs, here SnpEff (<http://snpeff.sourceforge.net/>), a tool for SNP annotation, was used to annotate ClinVar SNPs to genes of the human genome (Version GRCh38.86), to obtain Ensembl gene IDs. Then we mapped ClinVar genes to EPSD by using Ensembl gene IDs, and only SNPs directly changed known p-sites were reserved. For each record of ClinVar, columns entitled ‘RS_ID’, ‘Chrom’, ‘Pos’, ‘Ref’, ‘Alt’ and ‘Phenotype IDs’ were extracted for further integration.

*64) GWASdb (*[*http://jjwanglab.org/gwasdb*](http://jjwanglab.org/gwasdb)*) [**62**]*

We downloaded ‘gwasdb_20150819_snp_drug.gz’ from GWASdb (on February 5, 2019). ‘GENE_SYMBOL’ containing human gene names were used to map the data from GWASdb to EPSD. Finally, columns entitled ‘CHR’, ‘POS’, ‘SNPID’, ‘REF’, ‘ALT’, ‘P_VALUE’, ‘P_VALUE_TEXT’, ‘DURG_NAME’, ‘DRUG_ANNO’, ‘HPO_ID’ and ‘PMID’ were extracted for further integration.

*65) SNPdbe (*[*https://rostlab.org/services/snpdbe/*](https://rostlab.org/services/snpdbe/)*) [**63**]*

Single amino acid substitutions (SAASs) of non-synonymous single nucleotide polymorphisms (nsSNPs) with functional impacts annotated in the compressed file ‘SNPdbe_2012_03_05_sql’ were downloaded from the ‘Download’ page of SNPdbe (on May 15, 2019) [63]. From the sql files, 4 xml files including ‘seqs_refseq.xml’, ‘seqs_sp.xml’, ‘seqs_pmd.xml’ and ‘seqs_containingsnps.xml’ were extracted. The reference sequences and UniProt IDs of proteins were extracted from the files of ‘seqs_refseq.xml’ and ‘seqs_pmd.xml’, respectively. The positions and types of SAASs were extracted from the file ‘seqs_sp.xml’, as well as the corresponding species information, whereas associated functional impacts and diseases were retrieved from the file ‘geno2func.xml’. Peptides centred on SAAS residues flanked by 7 residues upstream and 7 residues downstream were retrieved. Then, UniProt IDs were used to find phosphorylated substrates in EPSD, while peptides were mapped to their corresponding protein sequences to pinpoint positions of SAASs. Only SAASs directly changed known p-sites were reserved.

*66) PMD (*[*http://pmd.ddbj.nig.ac.jp/~pmd/*](http://pmd.ddbj.nig.ac.jp/~pmd/)*) [**64**]*

Protein mutant data annotated in compressed files of ‘pmd.current’ and ‘pmdmuseq’ was downloaded in the ‘Download’ page of PMD (on May 2, 2019). The gene names and reference sequences of proteins were extracted from the file ‘pmdmuseq’, as well as the corresponding species information included in the lines started with ‘>’, while positions and types of SAAMs as well as ranges of long deletions were retrieved from the file ‘pmd.current’. The associated functional and structural impacts as well as diseases were also extracted. For SAAMs, peptides centred on point mutation residues flanked by 7 residues upstream and 7 residues downstream were retrieved, and mapped to EPSD. Only SAAMs directly changed known p-sites were reserved. For long deletions, only entries with at least one known p-site located in deletion regions were reserved. Here, obtained 30,161 protein SAAMs in 783 substrates.

*67) MSV3d (*[*http://decrypthon.igbmc.fr/msv3d/cgi-bin/home*](http://decrypthon.igbmc.fr/msv3d/cgi-bin/home)*) [**65**]*

Human missense variants mapped to 3D protein structures in the compressed file ‘allMut.txt.tar’ were downloaded from the ‘Home’ page of MSV3d (on May 14, 2019) [65]. The UniPort IDs were extracted from the first column, whereas positions and types of SAAMs were obtained from the second and third column, respectively. Associated functional impacts and diseases were obtained from the fourth column. Peptides centred on mutation residues flanked by 7 residues upstream and 7 residues downstream were retrieved. Then, UniProt IDs were used to find phosphorylated substrates in EPSD, while peptides were mapped to their corresponding protein sequences to pinpoint positions of SAAMs. Only SAAMs directly changed known p-sites were reserved. Here, we obtained 17,248 SAASs in 7,954 substrates.

*68) ActiveDriverDB (*[*https://activedriverdb.org/*](https://activedriverdb.org/)*) [**66**]*

ActiveDriverDB contained mutations that potentially influence post-translational modification (PTM) sites in human proteins/genes [66]. In this study, we downloaded ‘clinvar_mutations_affecting_ptm_sites.tsv’ and ‘mc3_mutations_affecting_ptm_sites.tsv’ from the ‘Download’ page of ActiveDriver (<https://activedriverdb.org/download/>, on Oct 14, 2022). Gene names were used to map data from ActiveDriverDB to EPSD. Information including ‘gene’, ‘muitation position’, ‘mutation alt’ and ‘mutation summary’ were retained in EPSD. Positions of mutations in the column ‘site position’ and ‘site residue’ were used, while only mutations directly changed known p-sites were reserved.

*69) BioMuta (*[*https://hive.biochemistry.gwu.edu/biomuta*](https://hive.biochemistry.gwu.edu/biomuta)*) [**67**]*

BioMuta contains human cancer-related sequence features from automated and manual curation and integration [67]. In this work, we downloaded ‘biomuta-master.csv’ and ‘biomuta-ac2genename.csv’ from BioMuta website (<https://hive.biochemistry.gwu.edu/beta/biomuta/content/BioMuta3.csv>, on May 21, 2024). UniProt IDs in file ‘biomuta-ac2genename.csv’ were used to map the data from BioMuta to EPSD, while columns including ‘ref_aa’, ‘alt_aa’, ‘chr_id’, ‘chr_pos’, ‘ref_nt’, ‘alt_nt’, ‘do_name’ and ‘pmid_list’, were integrated. Positions of mutations in the column ‘uniprot_pos’ were used, while only mutations directly changed known p-sites were reserved.

*70) Kin-Driver (*[*http://kin-driver.leloir.org.ar/*](http://kin-driver.leloir.org.ar/)*) [**68**]*

Kin-Driver is a database that provides information about the driver mutations of protein kinases.The sql file ‘kindriver_v82’ was downloaded from the ‘Download’ page of Kin-Driver (on May 21, 2024). From the sql file, 2 files ‘mutant’ and ‘disease’ were extracted. From the ‘mutant’ file, the UniProt IDs were extracted from the column ‘uniprot_id’, while positions and types of human SAAMs were extracted from the column ‘mutation’. The associated diseases were then extracted from the file ‘disease’. Then, SAAMs were mapped to known p-sites according to UniProt IDs.

*71) NECTAR (*[*http://nectarmutation.org/main/*](http://nectarmutation.org/main/)*) [**69**]*

The Non-synonymous Enriched Coding mutation Archive (NECTAR) is a data repository for the functionally important and disease-related amino acids. The file ‘disease_anno’ for human genes were downloaded from the FTP server of NECTAR (<ftp://ftp.nectarmutation.org/NECTAR/ProteinAnnotations/Diseases/>, on May 11, 2019). The Ensembl gene IDs were extracted from the column ‘ensp’, while original residues, mutant residues and positions of SAAMs were retrieved from the columns ‘p_ref’, ‘p_mut’ and ‘res_num’. The disease information in column ‘des’ was also considered. Then the mutations were mapped to known p-sites in *H. sapiens* according to Ensembl gene IDs and mutation positions.

*72) OMIM (*[*http://omim.org/*](http://omim.org/)*) [**70**]*

We downloaded ‘mim2gene.txt’ from OMIM (on May 21, 2024). Ensembl gene IDs were used to map human genes from OMIM to EPSD. Information in columns ‘MIM Number’, ‘MIM Entry Type’, ‘Entrez Gene ID’ and ‘Approved Gene Symbol’ were retained for further integration.

*73) PTMD (*[*http://ptmd.biocuckoo.org/*](http://ptmd.biocuckoo.org/)*) [**71**]*

Recently, we developed a well-curated database of PTMs that are associated with human Diseases (PTMD), containing 1,950 known PDAs in 749 proteins for 23 types of PTMs and 275 types of diseases curated from the literature. We downloaded the compressed file ‘PTM-Disease association.zip’ from the ‘DOWNLOAD’ page of PTMD. Columns including ‘Uniprot ID’, ‘Disease’, ‘PTMs Type’, ‘Position’ and ‘Literature’ were integrated, and the UniProt IDs were mapped to EPSD.

*74) MSDD (*[*http://www.bio-bigdata.com/msdd/*](http://www.bio-bigdata.com/msdd/)*) [**72**]*

The MiRNA SNP Disease Database (MSDD) documented experimentally supported associations among miRNAs, SNPs and human diseases [72]. We downloaded ‘msdd.txt’ from its website (<http://www.bio-bigdata.com/msdd/download.jsp>, on Oct 14, 2022). We extracted information, including ’Accession’, ‘PMID’, ‘miRNA’, ‘SNP’, ‘Disease’, ‘SNP position’, ‘Allele’, ‘Ancestral Allele’, ‘Method’, ‘Tissue/Cell line’, ‘Dysfunction Pattern’, ‘Population’, ‘Sample Size’, ‘Case-MAF’ and ‘Control-MAF’.

*75) DiseaseEnhancer (*[*http://biocc.hrbmu.edu.cn/DiseaseEnhancer/*](http://biocc.hrbmu.edu.cn/DiseaseEnhancer/)*) [**73**]*

DiseaseEnhancer provided a resource for human disease-associated enhancers [73]. We downloaded ‘enhInfo-1.0.2.txt’ and ‘enh2disease-1.0.2.txt’ from the download page of DiseaseEnhancer (http://biocc.hrbmu.edu.cn/DiseaseEnhancer/JumpToDownload, on May 21, 2024). Gene symbols were used to map data to EPSD. Columns entitled ‘id’, ‘VariantType’, ‘VariantName’, ‘Chr’, ‘Start’, ‘End’, ‘MutationType, ‘VariantConsequence’ and ‘PMID’ from the file ‘enhInfo-1.0.2.txt’ and column of ‘DiseaseType’ from the file ‘enh2disease-1.0.2.txt’ were extracted and integrated. Here, we obtained 348 human genes annotated with disease-associated enhancers.

**9. Protein-protein Interaction**

*76) IID (*[*http://iid.ophid.utoronto.ca/*](http://iid.ophid.utoronto.ca/)*) [**74**]*

All PPI files in IID database were downloaded from the ‘Download’ page of IID website ([*http://iid.ophid.utoronto.ca/*](http://iid.ophid.utoronto.ca/), on June 6, 2024), including ‘fly_annotated_PPIs.txt’, ‘human_annotated_PPIs.txt’, ‘mouse_annotated_PPIs.txt’, ‘rat_annotated_PPIs.txt’, ‘worm_annotated_PPIs’ and ‘yeast_annotated_PPIs.txt’. UniProt IDs were used to map the data from IID to EPSD. Information in columns including ‘symbol2’, ‘methods’, ‘pmids’, ‘dbs’, and ‘evidence type’ were retained for integration.

*77) iRefIndex (*[*http://irefindex.org*](http://irefindex.org)*) [**75**]*

We downloaded the file ‘All.mitab.22012018.txt.zip’ from iRefIndex website (<http://irefindex.org/download/irefindex/data/archive/release_15.0/psi_mitab/MITAB2.6/>, release 15, in Oct 14, 2022). UniProt IDs were used to map PPI pairs to EPSD. The information in columns including ‘method’, ‘pmids’, ‘taxa, ‘taxb’, ‘interactionType’, ‘sourcedb’, ‘confidence’, ‘edgetype’ and ‘numParticipants’ were integrated.

*78) PINA (*[*http://cbg.garvan.unsw.edu.au/pina*](http://cbg.garvan.unsw.edu.au/pina)*) [**76**]*First, we downloaded PPI files for 7 model species, including *A. thaliana* (‘Arabidopsis thaliana-20140521.tsv’), *C. elegans* (‘Caenorhabditis elegans-20140521.tsv’), *D. melanogaster* (‘Drosophila melanogaster-20140521.tsv’), *H. sapiens* (‘Homo sapiens-20140521.tsv’), *M. musculus* (‘Mus musculus-20140521.tsv’), *R. norvegicus* (‘Rattus norvegicus-20140521.tsv’) and *S. cerevisiae* (‘Saccharomyces cerevisiae-20140521.tsv’), from PINA website (<http://omics.bjcancer.org/pina/interactome.stat.do>, on Oct, 2022). UniProt IDs were used to map PPI pairs to EPSD. The information in columns entitled ‘Alt. ID(s) interactor A’, ‘Alt. ID(s) interactor B’, ‘Interaction detection method(s)’, ‘Publication Identifier(s)’, ‘Taxid interactor A’, ‘Taxid interactor B’, ‘Interaction type(s)’, ‘Source database(s)’ and ‘Interaction identifier(s)’ were integrated.

*79) HINT (*[*http://hint.yulab.org*](http://hint.yulab.org)*) [**77**]*

We downloaded all binary and co-complex PPI files from HINT website (<http://hint.yulab.org/download/>, on April 18, 2022). UniProt IDs were adopted to map PPI pairs to EPSD. We retained the information in columns including ‘Uniprot_A, ‘Uniprot_B’, ‘Gene_A’, ‘Gene_B’ and ‘pmid:method:quality’.

*80) Mentha (*[*http://mentha.uniroma2.it*](http://mentha.uniroma2.it)*) [**78**]*

We downloaded the file ‘all.zip’ from Mentha (<http://mentha.uniroma2.it/download.php>, Version 2019-05-13, on April 18, 2022). UniProt IDs were adopted as primary accessions to map the data to EPSD. We retained the information in columns including ‘Protein A’, ‘Gene A’, ‘Taxon A’, ‘Protein B’, ‘Gene B’, ‘Taxon B’, ‘Score’ and ‘PMID’.

*81) inBio Map^TM^ (*[*http://www.intomics.com/inbio/map*](http://www.intomics.com/inbio/map)*) [**79**]*

We downloaded the file ‘InBio_Map_core_2016_09_12.tar.gz’ from download page of inBio Map^TM^ (<https://www.intomics.com/inbio/map.html#downloads>, on April 18, 2022). UniProt IDs were adopted as primary accessions to map the data to EPSD. We retained the information in columns including ‘UP_ID’, ‘Gene_Name’, ‘Taxa_ID’, ‘MI_ID1’, ‘Interaction1’, ‘MI_ID2’, ‘Interaction2’ and ‘Confident score’.

*82) STRING (*[*https://string-db.org/*](https://string-db.org/)*) [**80**]*

We downloaded the file ‘protein.links.detailed.v12.0.txt.gz’ of 8 model organisms from the ‘Download’ page of STRING (Version 12, on June 6, 2024). Ensembl protein IDs were used to map PPI pairs to EPSD. The information in columns including ‘neighborhood’, ‘fusion’, ‘coocurence’, ‘coexpression’, ‘experimental’, ‘database’, ‘textmining’ and ‘combined_score’ were integrated.

*83) TIMBAL (*[*http://mordred.bioc.cam.ac.uk/timbal*](http://mordred.bioc.cam.ac.uk/timbal)*) [**81**]*

We downloaded ‘TIMBAL_sm.csv’ from TIMBAL website (http://mordred.bioc.cam.ac.uk /timbal/all, on May 18, 2019). UniProt IDs were used to map PPI pair to EPSD. Columns including ‘target_name’, ‘literature_name’, ‘bcomplex_descrip’, ‘pdb_code’, ‘activity_in_paper’, ‘assay_type’ and ‘pubmed_id’ were integrated.

**10. Drug-target relation**

84) TTD (<http://bidd.nus.edu.sg/group/cjttd/>) [82]

We downloaded the drug target information in raw format (‘P1-01-TTD_download.txt’) from the download page of TTD website (<https://db.idrblab.org/ttd/full-data-download>, last updated on September 12, 2017). UniProt IDs were used as primary accessions to find drug targeting proteins in EPSD. Drug names, type of targets, target validation, drug synonyms and associated diseases were extracted and integrated.

*85) DrugBank (*[*https://www.drugbank.ca/*](https://www.drugbank.ca/)*) [**83**]*

We downloaded the file ‘drugbank_all_full_database.xml.zip’ from the download page of DrugBank (<https://www.drugbank.ca/releases/latest>, version 2019-04-02, on May 19, 2019). UniProt IDs were used as primary accessions to find drug targeting proteins in EPSD. DrugBank IDs, drug names, drug type, groups, know-action and PMIDs were extracted and integrated.

*86) GtoPdb (*[*http://www.guidetopharmacology.org/targets.jsp*](http://www.guidetopharmacology.org/targets.jsp)*) [**84**]*

From GtoPdb (<http://www.guidetopharmacology.org/download.jsp>, on June 6, 2024), we downloaded the file ‘interactions.csv’. UniProt IDs were used as primary accessions to find drug targeting proteins in EPSD. The information in columns including ‘target’, ‘target_id’, ‘ligand’, ‘ligand_id’, ‘ligand_pubchem_sid’, ‘type’, ‘action’ and ‘pubmed_id’ were integrated.

*87) ADReCS-Target (*[*http://bioinf.xmu.edu.cn/ADReCS-Target*](http://bioinf.xmu.edu.cn/ADReCS-Target)*) [**85**]*

Adverse Drug Reaction Classification System-Target Profile (ADReCS-Target) is an update of DITOP and DART database. From ADReCS-Target (<http://bioinf.xmu.edu.cn/ADReCS-Target/download.jsp>, on May 20, 2024), we downloaded the file ‘P_D_A.xlsx’. UniProt IDs were used as primary accessions to find drug targeting proteins in EPSD. The information in columns including ‘BADD_TID’, ‘ADR_ID’, ‘ADReCS ID’, ‘ADR Term’, ‘Uniprot AC’ and ‘Drug_Name’ were integrated.

*88) ECOdrug (*[*http://www.ecodrug.org*](http://www.ecodrug.org)*) [**86**]*

The ECOdrug database is an open resource for the evolutionary conservation of human drug targets in various eukaryotic species [86]. From ECOdrug (http://www.ecodrug.org/#downloads, on May 19, 2019), we downloaded the file ‘ECOdrug_ensembl.csv’. We used Ensembl gene IDs as primary accessions to map the data from ECOdrug to EPSD. The information in columns entitled ‘Drug_name’, ‘MoA_text’, ‘DrugbankID’, ‘Drug_type’, ‘Drug_FirstApproval’, ‘ATC.code’, ‘Target_ChEMBLID’, ‘Target_pref_name’, ‘Target_name’, ‘Target_class’ and ‘interaction’ were integrated.

*89) DGIdb (*[*http://www.dgidb.org/*](http://www.dgidb.org/)*) [**87**]*

The drug-gene interaction database (DGIdb) provides the information of the druggable genome and drug-gene interactions [87]. We downloaded the file ‘interactions.tsv’ from DGIdb (<http://www.dgidb.org/downloads>, on May 20, 2024). We took NCBI Gene IDs as primary accessions to map drug-target relations to EPSD. We integrated the information in columns including ‘interaction_claim_source’, ‘interaction_types’, ‘drug_claim_name’, ‘drug_claim_primary_name’, ‘drug_name’ and ‘drug_chembl_id’.

*90) CTD (*[*http://ctdbase.org/*](http://ctdbase.org/)*) [**87**]*

The Comparative Toxicogenomics Database (CTD) is created for the purpose to investigate the relationships between environmental exposures and human health [87]. From CTD (<http://ctdbase.org/downloads/>, on May 20, 2024), we downloaded the file ‘CTD_chem_gene_ixns.tsv.gz’. NCBI Gene IDs were used as primary accessions to map drug-target relations to EPSD. The information in columns entitled ‘ChemicalName’, ‘ChemicalID’, ‘GeneSymbol’, ‘GeneForms, ‘Interaction’, ‘InteractionActions’ and ‘PubMedIDs’ were integrated.

*91) DrugCentral (*[*http://drugcentral.org/*](http://drugcentral.org/)*) [**88**]*

From DrugCentral (<http://drugcentral.org/download>, Postgres v14.5, in May 2024), the compressed file ‘drug.target.interaction.tsv’ was downloaded. Columns including ‘DRUG_NAME’ and ‘ACTION_TYPE’ were extracted, the UniProt IDs in the column ‘ACCESSION’ were mapped to EPSD.

**11. Orthologous information**

*92) InParanoid (*[*http://inparanoid.sbc.su.se/cgi-bin/index.cgi*](http://inparanoid.sbc.su.se/cgi-bin/index.cgi)*) [**89**]*

The compressed file ‘InParanoidUniProtXref’ was downloaded from the ‘Downloads’ page of InParanoid (Version 8.0, in May 2024). UniProt IDs were extracted as primary accessions to map the data to EPSD for the 8 model organisms.

*93) OMA (*[*https://omabrowser.org/oma/*](https://omabrowser.org/oma/)*) [**90**]*

The compressed file ‘oma-uniprot.txt’ was downloaded from the ‘Download’ page of OMA (Downloaded in May 2024). For each entry, the OMA ID was extracted. Then UniProt IDs were extracted and adopted as primary accessions to map the data to EPSD for the 8 model organisms.

*94) OrthoDB (*[*http://www.orthodb.org/*](http://www.orthodb.org/)*) [**91**]*

The compressed files ‘odb11v0_genes.tab’ and ‘odb11v0_OG2genes.tab’ were downloaded in the ‘Downloads’ page of OrthoDB (<https://www.orthodb.org/?page=filelist>, Version v11, in May 2023). The UniProt IDs and OrthoDB gene IDs in the column ‘Ortho DB unique gene id’ were extracted from the file ‘odb11v0_genes.tab’, while OrthoDB IDs were retrieved from the file ‘odb11v0_OG2genes.tab’. Then UniProt IDs were adopted as primary accessions to map the data to EPSD for the 8 model organisms.

*95) HOGENOM (*[*http://doua.prabi.fr/databases/hogenom/home.php?contents=query*](http://doua.prabi.fr/databases/hogenom/home.php?contents=query)*) [**92**]*

From the FTP server (<ftp://pbil.univ-lyon1.fr/pub/hogenom/release_06/>) of HOGENOM (Release 06, in May 2019), the file ‘expasy.HOGENOM’ was downloaded. For each entry, the gene family and UniProt ID were extracted from columns in ‘HOGENOM family’ and ‘Acc. Number in Uniprot’. Then UniProt IDs were adopted as primary accessions to map the data to EPSD for the 8 model organisms.

**12. Biological pathway**

*96) KEGG (*[*http://www.genome.jp/kegg/*](http://www.genome.jp/kegg/)*) [**93**]*

The files ‘pathway.list’, ‘genes_uniprot.list’ and ‘gene_map.tab’ for 8 model organisms were downloaded from the FTP server of Kyoto Encyclopedia of Genes and Genomes (KEGG) ([ftp.bioinformatics.jp/kegg](ftp://ftp.bioinformatics.jp/kegg), on Aug, 2021). For this study, we purchased a KEGG FTP subscription for personal use. UniProt IDs and associated pathways of proteins were extracted from the files ‘genes_uniprot.list’ and ‘gene_map.txt’, respectively. The detailed information of pathways was extracted from the file ‘pathway.list’. Then UniProt IDs were adopted as primary accessions to map the KEGG data to EPSD for the 8 model organisms.

*97) SignaLink (*[*http://signalink.org/*](http://signalink.org/)*) [**94**]*

The file ‘05032019-signalink-TuI8g3.csv’ was downloaded from the ‘download’ page of SignaLink (Version 3.0, on May 20, 2024). For each entry, the UniProt ID in the column ‘source_uniprotAC’ was extracted, as well as gene name, interaction type, directness and references. Then UniProt IDs were adopted as primary accessions to map the SignaLink data to EPSD for *H. sapiens*, *D. melanogaster* and *C. elegans*.

*98) Reactome (*[*https://reactome.org/*](https://reactome.org/)*) [**95**]*

The file ‘UniProt2Reactome.txt’ was downloaded from the ‘Download’ page of Reactome (Version 68, in May 2019). The UniProt IDs and pathway names were extracted. Then UniProt IDs were adopted as primary accessions to map the Reactome entries to EPSD for the 8 model organisms.

**13. Transcriptional regulator**

*99) TRRUST (*[*http://www.grnpedia.org/trrust/*](http://www.grnpedia.org/trrust/)*) [**96**]*

TRRUST is an expanded repository of transcriptional regulatory interactions focusing on human and mouse transcriptional regulatory networks [96]. ‘trrust_rawdata.human.tsv’ and ‘trrust_rawdata.mouse.tsv’ from download page of TRRUST (http://www.grnpedia.org /trrust/downloadnetwork.php, release 16, in April 2021) were downloaded. Gene names were used as primary keys to map data from TRRUST to EPSD. Columns entitled ‘T’, ‘Mode of Regulation’ and ‘References (PMID)’ were integrated.

*101) HEDD (*[*http://zdzlab.einstein.yu.edu/1/hedd/hedd.php*](http://zdzlab.einstein.yu.edu/1/hedd/hedd.php)*) [**97**]*

The Human Enhancer Disease Database (HEDD) contains comprehensive information of human enhancers and their association with human disease [97]. The files ‘DataDownload_Enhancer.txt’ and ‘DataDownload_EnhancerTagretGene.txt’ were downloaded from the ‘Download Data’ page of HEDD (<http://zdzlab.einstein.yu.edu/1/hedd/download.php>, on May 18, 2019). Gene names in the file ‘DataDownload_EnhancerTagretGene.txt’ were used to map data from HEDD to EPSD, information of enhancers in the file ‘DataDownload_Enhancer.txt’ were integrated.

*102) DroID (*[*http://droidb.org/*](http://droidb.org/)*) [**98**]*

The Drosophila Interactions Database (DroID) is a comprehensive interactions resource created for Drosophila. The file ‘tf_gene.txt’ was downloaded from the ‘Downloads’ page of DroID (<http://droidb.org/Downloads.jsp>, on May 18, 2024). Gene names in column ‘GENE_SYMBOL’ were adopted to map data from DroID to EPSD, information in columns entitled ‘TF_SYMBOL’, ‘PMID_METHOD’, ‘DATA_SOURCE_URL’ and ‘PUBMEDID’ were integrated.

*103) YTRP (*[*http://cosbi3.ee.ncku.edu.tw/YTRP/*](http://cosbi3.ee.ncku.edu.tw/YTRP/)*) [**99**]*

The Yeast Transcriptional Regulatory Pathway Database (YTRP) is constructed for yeast transcriptional regulatory pathways [99]. We downloaded the ‘TRP_direct_regulatory_network.txt’ from YTRP (http://cosbi3.ee.ncku.edu.tw/YTRP/Download, on May 18, 2019). Then, we used gene symbols as primary keys to map data from YTRP to EPSD. Columns entitled ‘TF’ and ‘Experimental Condition’ were integrated.

*104) RegNetwork (*[*http://www.regnetworkweb.org/*](http://www.regnetworkweb.org/)*) [**100**]*

The Regulatory Network Repository (RegNetwork) contains 5 type transcriptional and posttranscriptional regulatory relationships about human and mouse [100]. We downloaded the ‘human.zip’ and ‘mouse.zip’ from RegNetwork (http://www.regnetworkweb.org/download.jsp, on May 20, 2024). Then, we used Gene symbols in column ‘TARGET SYMBOL’ as primary keys to map data from RegNetwork to EPSD. We integrated information including ‘TARGET ID’, ‘REGULATOR SYMBOL’ and ‘REGULATOR ID’.

**14. mRNA expression**

*105) TCGA (*[*https://www.cancer.gov/tcga*](https://www.cancer.gov/tcga)*) [**24**]*

All available profiles of mRNA expression for 37 cancers, including ACC, BLCA, BRCA, CESC, CHOL, COAD, COADREAD, DLBC, ESCA, GBM, GBMLGG, HNSC, KICH, KIPAN, KIRC, KIRP, LAML, LGG, LIHC, LUAD, LUSC, MESO, OV, PAAD, PCPG, PRAD, READ, SARC, SKCM, STAD, STES, TGCT, THCA, THYM, UCEC, UCS, and UVM were downloaded from BROAD Institute (<http://gdac.broadinstitute.org/runs/stddata__2016_01_28/data>, Oncotated calls, level 3, in May 2022). In details, level-3 data such as ‘ACC.uncv2.mRNAseq_RSEM_normalized_log2_PARADIGM.txt’ were used to obtain mRNA expression level of each patient sample and Entrez IDs were used to map data from TCGA. Ultimately, ‘Project’, ‘Sample’, and ‘Expression’ were retained for integration.

*106) ICGC (*[*http://icgc.org/*](http://icgc.org/)*) [**25**]*

We downloaded all available gene expression profiles using sequencing-based platforms for 37 ICGC projects, including BLCA-US, BOCA-FR, BPLL-FR, BRCA-KR, BRCA-US, CESC-US, CLLE-ES, COAD-US, GBM-US, HNSC-US, KIRC-US, KIRP-US, LAML-US, LGG-US, LICA-FR, LIHC-US, LIRI-JP, LUAD-US, LUSC-US, MALY-DE, ORCA-IN, OV-AU, OV-US, PAAD-US, PACA-AU, PACA-CA, PAEN-AU, PBCA-US, PRAD-CA, PRAD-FR, PRAD-US, READ-US, RECA-EU, SKCM-US, STAD-US, THCA-US and UCEC-US, from the ICGC data portal (<https://dcc.icgc.org/releases/release_28/Projects/>, release 28, in March 2020). Ensembl gene IDs, Ensembl transcript IDs or gene symbols in the column ‘gene_id’ were mapped to EPSD. Columns, including ‘project_code’, ‘icgc_sample_id’, ‘normalized_read_count’ and ‘raw_read_count’, were extracted and integrated.

*107) ArrayExpress (*[*https://www.ebi.ac.uk/arrayexpress*](https://www.ebi.ac.uk/arrayexpress)*) [**101**]*

ArrayExpress provided microarray- and sequencing-based gene expression data [101]. We downloaded ‘allgenes_nonde_in_normal_2.0.14.tab’, ‘allgenes_nonde_in_organism_part_2.0.14.tab,’ ‘allgenes_updown_in_disease_2.0.19.tab’, ‘allgenes_updown_in_normal_2.0.14.tab’, ‘allgenes_updown_in_organism_part_2.0.14.tab’ and ‘organism_part_atlas_13.07.tab’ from ArrayExpress website (in June 2019). Ensembl gene IDs were used to map data to EPSD. We extracted information, including ‘Gene Name’, ‘Experimental Factor’, ‘Factor Value’, ‘Experiment Accession’, ‘Array Design Accession’, ‘Expression’ and ‘p Value’.

*108) GXD (*[*http://www.informatics.jax.org/expression.shtml*](http://www.informatics.jax.org/expression.shtml)*) [**102**]*

The Gene Expression Database (GXD) collected and integrated mouse developmental expression information. We downloaded both ‘MRK_GXDAssay.rpt.txt’ and ‘MGI_Gene_Model_Coord.rpt.txt’ from its website (http://www.informatics.jax.org/mgihome/GXD/aboutGXD.shtml, in May 2024). Ensembl gene IDs were used to map data from GXD to EPSD. We extracted information including ‘Marker ID’, ‘Marker Symbol’, ‘Marker Name’ and ‘MGI Assay Accession ID’.

*109) COSMIC (*[*https://cancer.sanger.ac.uk/cosmic*](https://cancer.sanger.ac.uk/cosmic)*) [**26**]*

The compressed file ‘CosmicCompleteGeneExpression.tsv’ was downloaded from the ‘Downloads’ page of COSMIC (<https://cancer.sanger.ac.uk/cosmic/download>, release v98, on June, 2023). Columns including ‘SAMPLE_NAME’, ‘REGULATION’, ‘Z_SCORE’ and ‘ID_STUDY’ were integrated, and gene symbols in the column ‘GENE_NAME’ were used to map data to EPSD.

*110) BioXpress (*[*https://hive.biochemistry.gwu.edu/bioxpress*](https://hive.biochemistry.gwu.edu/bioxpress)*) [**67**]*

The BioXpress database provides the gene/miRNA expression profiled from cancer samples. We download the file ‘BioXpress_gene_differential_expression_v2.0.csv’ from its website (Version 2.0, in May 2019). Columns, including ‘UniProtKB_AC’, ‘log2FoldChange’, ‘p_value’, ‘Significant’, ’Trend’, ‘TCGA Cancer’, ‘Cancer Ontology’, ‘#Patients’ and ‘UBERON_ID’ were integrated, and UniProt IDs were mapped to EPSD.

*111) TissGDB (*[*http://zhaobioinfo.org/TissGDB*](http://zhaobioinfo.org/TissGDB)*) [**103**]*

The Tissue specific Gene DataBase in cancer (TissGDB) is a tissue-specific gene annotation database for cancer [103]. First, the ‘Tissg_DEGs’ and ‘TissgDB_basic_uniq.txt’ files were download from the ‘Download’ page of TissGDB (<https://bioinfo.uth.edu/TissGDB/download.html>, in May 2019). Then, columns including ‘Cancer Type’, ‘Mean Expression in Tumor’, ‘Mean Expression in Normal’, ‘-log2(FC)’, ‘Pvalue’ and ‘FDR’ were integrated from ‘Tissg_DEGs’, and gene symbols were obtained in the column ‘TissGene’. Finally, UniProt IDs and gene symbols were extracted from ‘TissgDB_basic_uniq.txt’, and UniProt IDs were mapped to EPSD.

*112) FFGED (*[*http://bioinfo.townsend.yale.edu/*](http://bioinfo.townsend.yale.edu/)*) [**104**]*

The filamentous fungal gene expression database (FFGED) provided comprehensive information for gene expression in filamentous fungal [104]. All 13,420 XML profiles were downloaded from the FTP server of FFGED. Ensembl gene IDs were used to map data to EPSD. Columns entitled ‘Experiment ID’, ‘Experiment Name’, ‘Expression Variable’ and ‘Expression Value’ were integrated.

*113) SZDB (*[*http://www.szdb.org/*](http://www.szdb.org/)*) [**105**]*

The schizophrenia database (SZDB) is a comprehensive resource for Schizophrenia, and related gene expression data was integrated [105]. 9 files including ‘clusters.txt’, ‘entries.txt’, ‘gene.txt’, ‘MD_CBC.txt’, ‘PFC_MSC.txt’, ‘Score.csv’, ‘stages.txt’, ‘STR_AMY.txt’ and ‘V1C_STC.txt’ were downloaded from ‘Download’ page of SZDB (http://www.szdb.org/download.html, in February 2018). Gene symbols were used to map data to EPSD. We retained columns including ‘Gene1 ID’, ‘Gene1’, ‘Pearson’, ‘Stage’ and ‘Cluster’ for further integration.

*114) TISSUES (*[*http://tissues.jensenlab.org/*](http://tissues.jensenlab.org/)*) [**106**]*

The TISSUES integrates the knowledge on tissue expression levels of mRNAs manually curated from the literature, proteomic and transcriptomic profilings, as well as automatic text mining [106]. We downloaded all knowledge, experiments and text mining files of 3 model organisms, including *H. sapiens*, *M. musculus* and *R. norvegicus*, from the ‘Downloads’ page of TISSUES (<https://tissues.jensenlab.org/Downloads>, Version 2.0, in June 2024). Then, ‘source database’ and ‘evidence type’ columns of knowledge files, ‘source dataset’ and ‘expression score’ columns of experiments files, and ‘z-score’ column of text mining files were extracted, respectively. For all files, columns including ‘tissue identifier’, ‘tissue name’ and ‘confidence score’ were integrated, and Ensembl protein IDs were mapped to EPSD.

*115) The Human Protein Atlas (*[*http://www.proteinatlas.org/*](http://www.proteinatlas.org/)*) [**107**]*

The Human Protein Atlas (HPA) provides a map of human proteins in cells, tissues and organs with various omics technologies, including antibody-based imaging, MS-based proteomics, transcriptomics and systems biology [107]. Here, only human transcriptomic data was considered, and we downloaded all compressed files, including ‘rna_celline.tsv’ and ‘rna_tissue.tsv’, for 64 cell lines and 37 tissues based on RNA-seq from the ‘DOWNLOADABLE DATA’ page of HPA (<https://www.proteinatlas.org/about/download>, Version 23.0, in May 2024). Then, all samples in ‘Sample’ column and transcripts per million in ‘Value’ column were integrated, and Ensembl gene IDs in the column ‘Gene’ were mapped to EPSD.

*116) Human Proteome Map (*[*http://www.humanproteomemap.org/*](http://www.humanproteomemap.org/)*) [**108**]*

The Human Proteome Map (HPM) portal is an interactive resource of human proteome [108]. Here, only human transcriptomic data integrated in HPM was considered. The file ‘HPM_gene_level_epxression_matrix_Kim_et_al_052914.csv’ was downloaded from the ‘Download’ page of HPM (<http://www.humanproteomemap.org/download_hpm_data.php>, in May 2024). Then, gene symbols in the column ‘Gene’ were mapped to EPSD. We integrated the mRNA expression information of 17 adult tissues, 6 primary hematopoietic cells and 7 fetal tissues, including ‘Fetal Heart’, ‘Fetal Liver’, ‘Fetal Gut’, ‘Fetal Ovary’, ‘Fetal Testis’, ‘Fetal Brain’, ‘Adult Frontal Cortex’, ‘Adult Spinal Cord’, ‘Adult Retina’, ‘Adult Heart’, ‘Adult Liver’, ‘Adult Ovary’, ‘Adult Testis’, ‘Adult Lung’, ‘Adult Adrenal’, ‘Adult Gallbladder’, ‘Adult Pancreas’, ‘Adult Kidney’, ‘Adult Esophagus’, ‘Adult Colon’, ‘Adult Rectum’, ‘Adult Urinary Bladder’, ‘Adult Prostate’, ‘Placenta’, ‘B Cells’, ‘CD4 Cells’, ‘CD8 Cells’, ‘NK Cells’, ‘Monocytes’ and ‘Platelets’.

**15. Protein expression/Proteomics**

*117) The Human Protein Atlas (*[*http://www.proteinatlas.org/*](http://www.proteinatlas.org/)*) [**107**]*

The compressed file ‘proteinatlas.xml’ was downloaded from the ‘DOWNLOADABLE DATA’ page of HPA (<https://www.proteinatlas.org/about/download>, Version 23.0, in May 2024). Here, the protein expression data was integrated. Scores of protein expression except ‘Not detected’ with corresponding tissues were extracted, and the UniProt IDs were mapped to EPSD. Finally, the protein expression of 44 tissues, including ‘appendix’, ‘breast’, ‘bronchus’, ‘cerebral cortex’, ‘cervix, uterine’, ‘colon’, ‘duodenum’, ‘endometrium’, ‘epididymis’, ‘esophagus’, ‘fallopian tube’, ‘gallbladder’, ‘kidney’, ‘liver’, ‘lung’, ‘nasopharynx’, ‘oral mucosa’, ‘pancreas’, ‘parathyroid gland’, ‘placenta’, ‘prostate’, ‘rectum’, ‘salivary gland’, ‘seminal vesicle’, ‘skin’, ‘small intestine’, ‘stomach’, ‘testis’, ‘thyroid gland’, ‘tonsil’, ‘urinary bladder’, ‘vagina’, ‘adrenal gland’, ‘bone marrow’, ‘caudate’, ‘cerebellum’, ‘heart muscle’, ‘hippocampus’, ‘lymph node’, ‘ovary’, ‘skeletal muscle’, ‘smooth muscle’, ‘soft tissue’ and ‘spleen’ were curated.

*118) Human Proteome Map (*[*http://www.humanproteomemap.org/*](http://www.humanproteomemap.org/)*) [**108**]*

The file ‘HPM_protein_level_expression_matrix_Kim_et_al_052914.csv’ was downloaded from the ‘Download’ page of HPM (http://www.humanproteomemap.org/download_hpm_data.php). The protein expression of 17 adult tissues, 6 primary hematopoietic cells and 7 fetal tissues, including ‘Fetal Heart’, ‘Fetal Liver’, ‘Fetal Gut’, ‘Fetal Ovary’, ‘Fetal Testis’, ‘Fetal Brain’, ‘Adult Frontal Cortex’, ‘Adult Spinal Cord’, ‘Adult Retina’, ‘Adult Heart’, ‘Adult Liver’, ‘Adult Ovary’, ‘Adult Testis’, ‘Adult Lung’, ‘Adult Adrenal’, ‘Adult Gallbladder’, ‘Adult Pancreas’, ‘Adult Kidney’, ‘Adult Esophagus’, ‘Adult Colon’, ‘Adult Rectum’, ‘Adult Urinary Bladder’, ‘Adult Prostate’, ‘Placenta’, ‘B Cells’, ‘CD4 Cells’, ‘CD8 Cells’, ‘NK Cells’, ‘Monocytes’ and ‘Platelets’ were integrated, and RefSeq protein IDs in the column ‘RefSeq Accession’ were used to map data to EPSD.

**16. Subcellular localization**

*119) NLSdb (*[*https://rostlab.org/services/nlsdb/*](https://rostlab.org/services/nlsdb/)*) [**109**]*

The NLSdb annotated nuclear export and localization signals (NES/NLS) of proteins [109]. The ‘signals.csv’ and ‘extsignals.csv’ files were downloaded from the ‘Downloads’ page of NLSdb (<https://rostlab.org/services/nlsdb/downloads>, in June 2024). Columns including ‘Sequence’, ‘SignalType’, ‘AnnotationType’, ‘ConfidenceNuc’ and ‘ConfidenceFam’ were integrated, and the UniProt IDs in the column ‘Origin’ were mapped to EPSD.

*120) COMPARTMENTS (*[*https://compartments.jensenlab.org/*](https://compartments.jensenlab.org/)*) [**110**]*

The COMPARTMENTS database integrated the knowledge of protein subcellular localizations [110]. We downloaded all .tsv files of channels integrated annotation in 7 model organisms, including *H. sapiens*, *M. musculus*, *R. norvegicus*, *D. melanogaster*, *C. elegans*, *A. thaliana* and *S. cerevisiae*, from the ‘Downloads’ page of COMPARTMENTS (<https://compartments.jensenlab.org/Downloads>, in June 2024). Columns including ‘Name’ and ‘Condifence’ were extracted, and Ensembl protein IDs were mapped to EPSD.

*121) Translocatome (*[*http://translocatome.linkgroup.hu*](http://translocatome.linkgroup.hu)*) [**111**]*

The Translocatome database provides annotation of translocating proteins of human cells, by collection, manual curation and prediction [111]. The ‘translocatome_all_data.csv’, ‘translocatome_transloc_data.csv’ and ‘translocatome_nontransloc_data.csv’ files were downloaded from the ‘DOWNLOAD’ page of Translocatome (<http://translocatome.linkgroup.hu/download>, in June 2024). Then, the UniProt IDs were extracted and mapped to EPSD with corresponding ‘translocation evidence score’ column from ‘translocatome_all_data.csv’. Finally, manually curated translocating and non-translocating proteins were extracted from ‘translocatome_transloc_data.csv’ and ‘translocatome_nontransloc_data.csv’, respectively.

**Supplementary Tables**

**Supplementary Table S1.** A summary of mainstream p-site databases for eukaryotic phosphorylation.

**Supplementary Table S2**. The public data resources that included 10 phosphorylation databases and 100 additional databases or tools.

**Supplementary Table S3**. The distribution of p-sites and phosphoproteins across the 223 species.

**Supplementary Table S4**. The effect types of 88,074 functional events.

**Supplementary Table S5**. The comparison of EPSD 2.0 and EPSD 1.0.
