## Supplementary figures and images for "EPSD 2.0: An Updated Database of Protein Phosphorylation Sites across Eukaryotic Species"

### Supplementary Figure S1.tiff

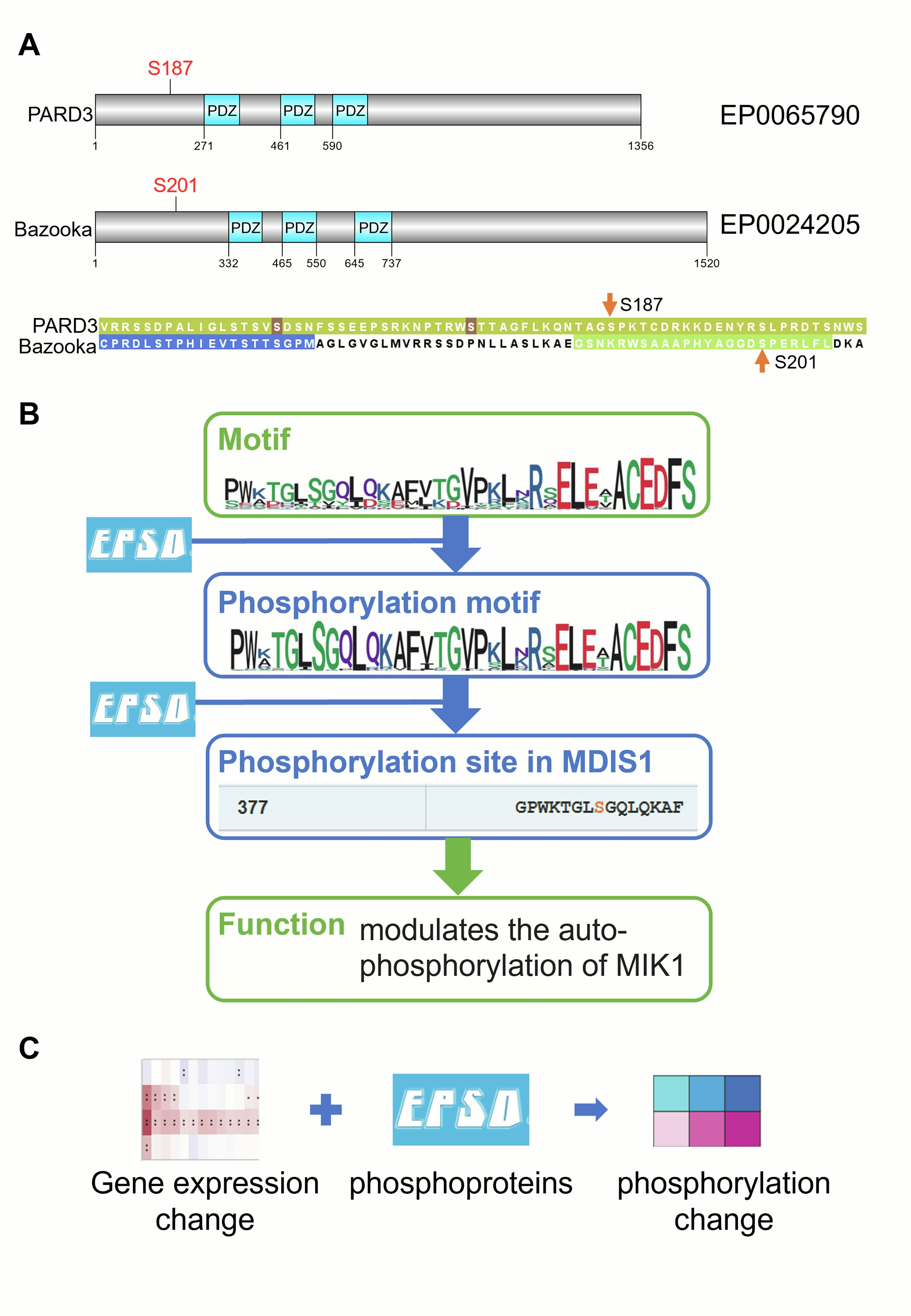
